# Genetic variation in the activity of a TREM2-p53 signaling axis determines oxygen-induced lung injury

**DOI:** 10.1101/2024.09.13.612775

**Authors:** Yohei Abe, Nathaneal J. Spann, Wenxi Tang, Fenghua Zeng, John Lalith Charles Richard, Cadence Seymour, Sean Jansky, Miguel Mooney, Robert Huff, Kelly Chanthavixay, Debanjan Dhar, Souradipta Ganguly, Jason L. Guo, David M. Lopez, Michael T. Longaker, Christopher Benner, Christopher K. Glass, Eniko Sajti

**Affiliations:** Department of Cellular and Molecular Medicine, University of California San Diego, La Jolla, CA 92093, USA; Department of Medicine, University of California San Diego, La Jolla, CA 92093, USA; Department of Pediatrics, University of California San Diego, California, La Jolla, CA 92093, USA; Hagey Laboratory for Pediatric Regenerative Medicine, Division of Plastic and Reconstructive Surgery, Stanford University School of Medicine, Stanford, CA 94305, USA; Department of Surgery, Stanford University School of Medicine, Stanford CA 94305, USA; Institute for Stem Cell Biology and Regenerative Medicine, Stanford University School of Medicine, Stanford, CA 94305, USA; Department of Medicine, Division of Endocrinology, University of California San Diego, La Jolla, CA 92093, USA

## Abstract

Bronchopulmonary dysplasia (BPD), a chronic lung disease, is the most common major complication of preterm birth. Supplemental oxygen administration, while lifesaving in the neonatal period, remains a key determinant of BPD pathophysiology. Exposure of the immature lung to increased levels of oxygen elicits an inflammatory response resulting in abnormal lung development. However, not every premature infant is equally sensitive to develop BPD. Using genetically diverse mouse strains, we show that the innate immune response activated in the lungs of mice sensitive to hyperoxia that develop BPD-like lung injury differs from mice resilient to disease. Specifically, we identified a selective upregulation of triggering receptor expressed on myeloid cells 2 (TREM2) on lung macrophages and monocytes in the hyperoxia-sensitive C57BL/6J mouse strain. We show that loss of function of TREM2 signaling in myeloid cells resulted in a dramatically improved phenotype after neonatal hyperoxia exposure characterized by a dampened immune response, preserved alveolar structure, and preserved cell proliferative potential supporting normal lung development. At the molecular level, inhibition of TREM2 signaling dampened the magnitude of p53 activation and resulted in cell cycle arrest instead of apoptosis. These findings show that TREM2 is a critical regulator of the pathogenic innate immune response to hyperoxia and highlight its importance as a potential therapeutic target for mitigating injury in the hyperoxia-exposed developing lung.

## Introduction

Not every premature infant is equally susceptible to bronchopulmonary dysplasia (BPD), the chronic lung disease associated with premature birth ^1, 2, 3^. BPD primarily affects premature infants who have been exposed to mechanical ventilation and oxygen therapy. Despite exposure to very similar therapeutic interventions, the progress of infants through the neonatal intensive care unit will vary widely, some developing mild symptoms while others developing severe respiratory complications. This implies that factors inherent to the premature infant modulate the capacity of the developing lung to respond to the life-sustaining therapies and thereby determine long-term outcomes.

Oxygen therapy, while lifesaving, has significant side effects. Exposure of the developing lung to increased oxygen concentration, also called hyperoxia, injures the developing lung. At a cellular level, hyperoxic lung injury is characterized by alveolar cell apoptosis and necrosis ^4^ that is accompanied by an inflammatory response. In susceptible individuals, the injury permanently derails normal lung development leading to structural and functional changes that can last life-long ^5, 6^. However, it is unclear how the processes of cell death combine with inflammation to produce a variable degree of injury and regenerative potential resulting in the different lung phenotypes observed in patients with BPD.

Although inflammation is a key driver in the pathogenesis of BPD, the molecular triggers, sensory receptors and signaling pathways of the immune cells that respond to the hyperoxia-induced injury are incompletely understood. Polymorphisms affecting the immune response were found to be predisposing to BPD ^7^, therefore further exploration of the genetic basis of the immune response is warranted. Over the last few years, it has become clear that the pathogenic processes by which genetic variants influence disease are cell type specific ^8^. In addition, ∼80-90% of the common risk variants for disease susceptibility reside in non-coding regions of the genome. Therefore, understanding how genetic variation in immune cells contributes to susceptibility or resilience against injury in the developing lung has the potential to uncover new targets for immunomodulation.

Mouse strains exhibit genetic variations that influence their susceptibility to various diseases and can be used to study individual differences in disease susceptibility ^9, 10, 11, 12^. We have chosen two strains with known susceptibility or resistance to hyperoxia-induced lung injury to study the genetic factors that contribute to the severity of neonatal hyperoxia-induced injury. C57BL/6J mice have been shown to be sensitive to hyperoxia-induced lung injury, while DBA/2J mice are resistant ^13^. These two mouse strains exhibit significant genetic differences in multiple lung myeloid cell subsets including macrophages and monocytes ^14^. These differences may contribute to variations in susceptibility to hyperoxia-induced lung injury between the two strains making them a great choice for studying specific aspects of myeloid cell biology in hyperoxic lung injury.

The lung is populated by a wide variety of immune cells ^15^. In case of injury, the first line of cellular defense is provided by innate immune cells including lung macrophages and monocytes. In addition, myeloid cells play an important role in lung development and homeostasis. After hyperoxia exposure, resident myeloid cells are activated and recruited neutrophils and monocytes from the blood further add to the myeloid cell diversity in the injured lung ^16, 17^. All these cells can contribute to the pathogenesis of BPD through their involvement in inflammation, oxidative stress, tissue remodeling, and immune regulation. However, the molecular pathways and factors which determine whether the immune response is protective, or damaging are not known. Triggering receptor expressed on myeloid cells 2 (TREM2) is a protein receptor selectively expressed on the surface of myeloid cells including macrophages and monocytes. It has garnered significant attention in recent years due to its crucial role in modulating the immune response, particularly in the context of neuroinflammation and neurodegenerative diseases ^18, 19^. While research on TREM2 in lung diseases is still in its early stages, the available evidence suggests that this receptor plays a role in regulating immune responses, inflammation, and tissue repair in the lung ^20, 21, 22, 23^.

Here we show that TREM2 plays a central role in orchestrating the myeloid response to hyperoxia exposure in the developing lung. Exploiting natural genetic variation between hyperoxia-sensitive (C57BL/6J) and resistant (DBA/2J) strains of mice, we identified a selective upregulation of TREM2 on lung macrophages and monocytes in the hyperoxia-sensitive C57BL6/J strain. Importantly, genetic ablation of *Trem2* rendered hyperoxia-sensitive mice resistant to injury. TREM2-deficient mice had a dramatically improved phenotype after neonatal hyperoxia exposure characterized by a dampened immune response, preserved alveolar structure, and preserved cell proliferative potential supporting normal lung development.

These findings show that TREM2 serves as a critical regulator of the immune response to hyperoxia and highlight its importance as a potential therapeutic target for mitigating injury in the hyperoxia exposed developing lung.

## Results

### Differential activation of apoptosis and cell proliferation underlies strain differences in the severity on neonatal hyperoxia-induced lung injury

In various mouse strains, there is a significant difference in susceptibility to hyperoxia-induced lung injury, suggesting that response to oxygen is at least, in part, influenced by genetic background. When exposed to high concentrations of oxygen, DBA/2J (DBA) mice show significantly less lung injury as compared to C57BL/6J (B6) mice ^13, 24, 25^. To test the hypothesis that a comparison between hyperoxia sensitive B6 and hyperoxia resistant DBA mice will identify gene programs that promote resistance to lung injury, we exposed newborn mice from both strains to 75% oxygen for 14 days (widely used model for bronchopulmonary dysplasia (BPD)) (Fig. 1a, left panel). In both strains, hyperoxia exposure resulted in similar weight loss (Fig. 1a, right panel), eliminating differences in weight gain as a contributor to differences in organ development. Analyzing the hyperoxia-induced lung injury in the two strains, we observed significant differences in lung morphology. While hyperoxia exposure led to severe alveolar simplification in B6 mice, lung tissue in DBA mice was preserved (Fig. 1b). To quantify these changes, we measured the mean linear intercept and observed a significant increase in alveolar diameter in hyperoxia-exposed B6 mice, while alveolar architecture did not show significant changes in the resistant DBA strain (Fig. 1b).

**Fig. 1.**
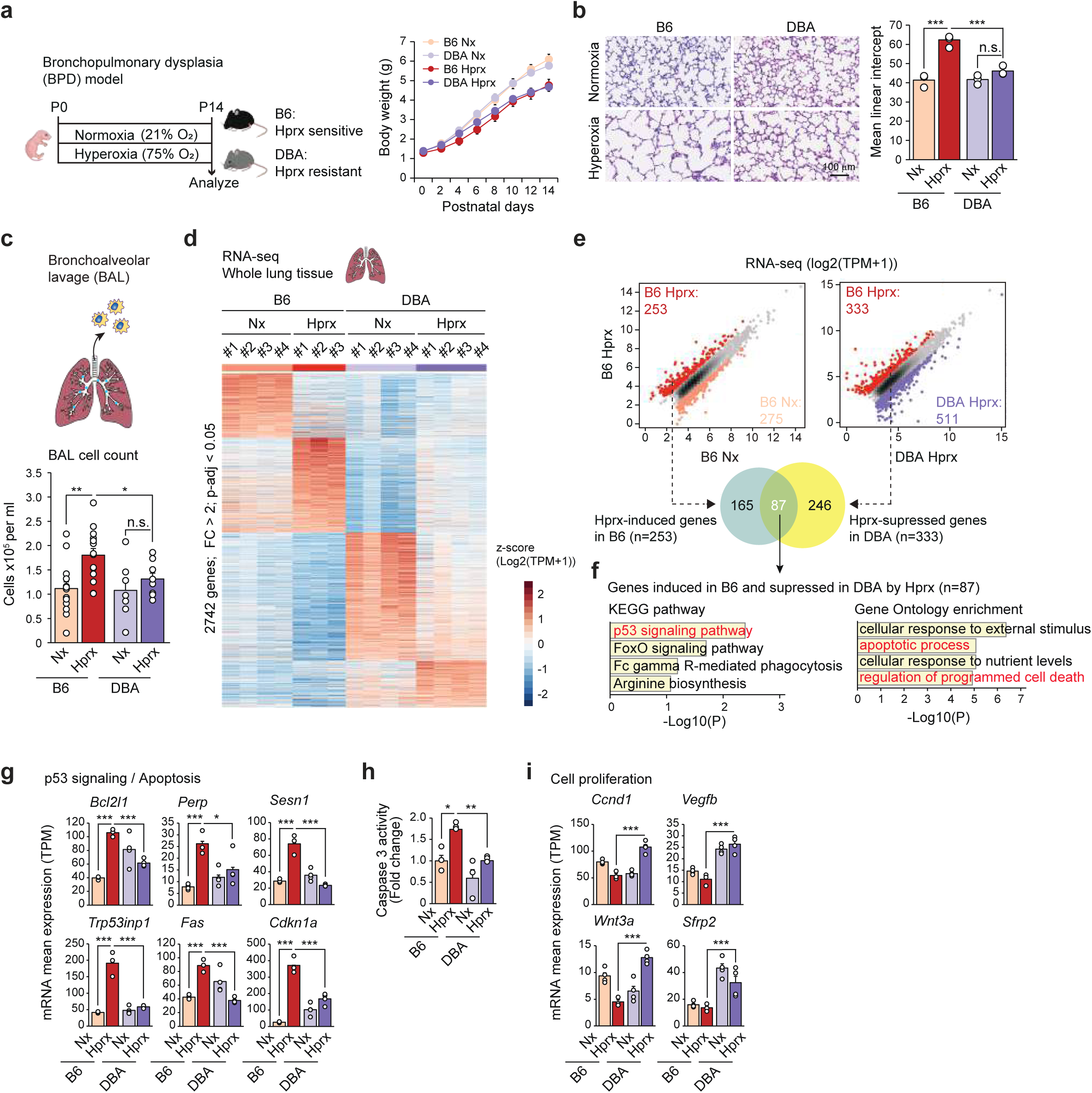
Strain differences in neonatal hyperoxia-induced lung injury. (a) Neonatal C57BL/6J (B6) and DBA/2J (DBA) mice were exposed to 75% oxygen (Hyperoxia) from P0 to P14 and harvested at P14 for analysis. Littermate controls raised in room air (21% oxygen, Normoxia) were served as controls. Body weights in hyperoxia-exposed B6 and DBA mice. Data are mean ± SEM (n=6 for each group). (b) H&E staining was performed on formalin fixed paraffin embedded lungs to assess the alveolar complexity at P14 in B6 mice (left panels) and DBA mice (right panels) with scale bars denoting 100 μm. The bar graph represents the results of the quantification of alveolar simplification using mean linear intercept. Data are mean ± SEM (n=3-4 for each group). ANOVA was performed followed by Tukey’s post hoc comparison. \*\*\**p* < 0.001. n.s.: not significant. (c) Bronchoalveolar lavage (BAL) was performed at P14 in hyperoxia-exposed B6 and DBA mice, and the control mice. Data are mean ± SEM (n=14 for B6 and n=7-8 for DBA). ANOVA was performed followed by Tukey’s post hoc comparison. \**p* < 0.05 and \*\**p* < 0.01. n.s.: not significant. (d) RNA-seq in whole lung tissues of B6 and DBA mice at P14. Heatmap shows the 2742 differentially expressed genes (FC > 2; *p*-adj < 0.05) comparing hyperoxia-exposed B6 and DBA mice, and the normoxic controls. (e) Scatterplots of RNA-seq data in whole lung tissues showing hyperoxia-regulated gene expression in B6 mice (left panel) and DBA mice-regulated gene expression in the hyperoxic conditions (right panel). Dark red dots in left panel: significant hyperoxia-induced genes in B6 mice, light red dots in left panel: significant hyperoxia-suppressed genes in B6 mice, dark red dots in right panel: significant DBA mice-suppressed genes in the hyperoxic conditions, dark purple dots in right panel: significant DBA mice-induced genes in the hyperoxic conditions (FC > 2; FDR < 0.05). Venn diagram shows the overlap between hyperoxia-induced genes in B6 mice (n = 253) and hyperoxia-suppressed genes in DBA mice (n = 333). (f) KEGG pathway (left panel) and Gene Ontology enrichment (right panel) analysis of the 87 genes that were induced in B6 mice and suppressed in DBA mice by hyperoxia. (g) Bar plots for expression of representative genes belonging to the p53 signaling pathway. Data are mean ± SEM. \**p*-adj < 0.05 and \*\*\**p*-adj < 0.001. (h) Caspase 3 activity measured in cytosolic fractions from whole lung tissues (B6 and DBA mice) harvested on P14. Data are mean ± SEM (n=3 for each group). ANOVA was performed followed by Tukey’s post hoc comparison. \**p* < 0.05 and \*\**p* < 0.01. (i) Bar plots for expression of representative genes belonging to the cell proliferation pathway. Data are mean ± SEM. \*\*\**p*-adj < 0.001. Each dot represents one mouse (b, c, g, h, i). See also Extended Data Fig. 1.

As lung injury elicits an inflammatory response, we assessed the severity of the inflammation caused by hyperoxia, by performing bronchoalveolar lavage (BAL) at the end of the exposure on postnatal day (PND) 14. We observed an increase in the number of immune cells in the BAL fluid of hyperoxia exposed B6 mice but not in the BAL fluid of DBA mice (Fig. 1c), implying a dampened immune response and less inflammatory cell recruitment to the airways of hyperoxia resistant DBA mice. Furthermore, these findings suggest an association between the severity of inflammatory response and impaired lung development.

To elucidate the molecular mechanisms directing resilience against hyperoxia-induced injury, we performed RNA sequencing (RNA-seq) in the whole lung and measured differences in gene expression between the two strains (Fig. 1d). We detected striking differences in the hyperoxia-induced gene programs between B6 and DBA mice. Of the 253 genes induced > 2-fold by hyperoxia (p-adj < 0.05) in B6 mice, 87 genes were repressed in hyperoxia-exposed DBA mice relative to hyperoxia-exposed C57 mice (Fig. 1e, Venn diagram). These 87 genes induced in B6 by hyperoxia but repressed in DBA mice were enriched in genes related to the p53 signaling pathway, apoptosis and cell death (Fig. 1f). For example, *Bcl2l1* (encoding a pro-apoptotic regulator that controls the production of reactive oxygen species in mitochondria), *Prep* (encoding a cytosolic prolyl endopeptidase that cleaves peptide bonds on the C-terminal side of prolyl residues within peptides), *Sesn1* (encoding a member of the sestrin family known to mediate the cellular response to oxidative stress), *Trp53inp1* (encoding a tumor protein 53-induced nuclear protein that modulates p53 transcriptional activity), *Fas* (encoding a member of the TNF-receptor superfamily that plays a central role in the physiological regulation of programmed cell death), and *Cdkn1a* (encoding a protein that functions as a cyclin-dependent kinase inhibitor and also known as p21) were induced by hyperoxia in B6 mice but expressed at much lower levels in DBA mice (Fig. 1g). Consistent with the apoptotic gene expression, hyperoxia exposure resulted in significantly higher caspase 3 activity in cytosolic fractions from whole lungs of the hyperoxia-sensitive B6 but not the hyperoxia-resistant DBA mice (Fig. 1h). Importantly, 95 genes induced by hyperoxia selectively in the DBA strain were related to cell proliferation referred to as cell cycle and cell division (Extended Data Fig. 1a), exemplified by *Ccnd1* (encoding a cyclin family member), *Vegfb* (encoding a protein involved in vasculogenesis), *Wnt3a* (encoding a protein regulating cell fate and patterning), and *Sfrp2* (encoding a soluble modulator of Wnt signaling) (Fig. 1i). Baseline differences in gene expression between the two strains are shown in Extended Data Fig. 1b. The subset of higher expressed genes in B6 mice (n=538) did not indicate apoptotic genes, suggesting the expression of genes related to apoptosis and cell death depends on hyperoxia exposure.

Taken together, these data demonstrate strain specific responses to neonatal hyperoxia exposure and identify differential activation of apoptosis- and cell proliferation-related gene programs as modulators of the severity of lung injury.

### Trem2 deletion in myeloid cells protects the developing lung from neonatal hyperoxia-induced injury

Hyperoxia exposure leads to inflammation in the lung and an excessive immune response can further exacerbate the lung injury by tipping the balance between cell survival and cell death pathways. To identify hyperoxia-induced gene programs specific to lung macrophages and monocytes, we analyzed the genes induced in alveolar macrophages, interstitial macrophages and classical monocytes in a previously published single-cell RNA-seq data set ^16^. While hyperoxia induced cell-specific gene programs in the two lung macrophage subsets and monocytes (Extended Data Fig. 2a, b, c), we identified 18 genes that were commonly induced by hyperoxia in all three myeloid cell subsets (Extended Data Fig. 2d). Of these 18 genes, only one gene, *Trem2* was detected in the whole lung RNA-seq data (Fig. 1e and Extended Data Fig. 2d). To gain further insight in the cell type specific regulation of *Trem2*, we compared gene expression levels in hyperoxia-exposed and control lungs at single cell level (Fig. 2a). As expected, *Trem2* expression was restricted to myeloid cells, with interstitial macrophages expressing the highest level at baseline. The *Trem2* gene was also expressed in alveolar macrophages and classical monocytes but absent in other myeloid cell subsets such as neutrophils and dendritic cells. Importantly, hyperoxia exposure resulted in *Trem2* induction in all three myeloid cell subsets (alveolar macrophages, interstitial macrophages, classical monocytes) (Fig. 2a).

**Fig. 2.**
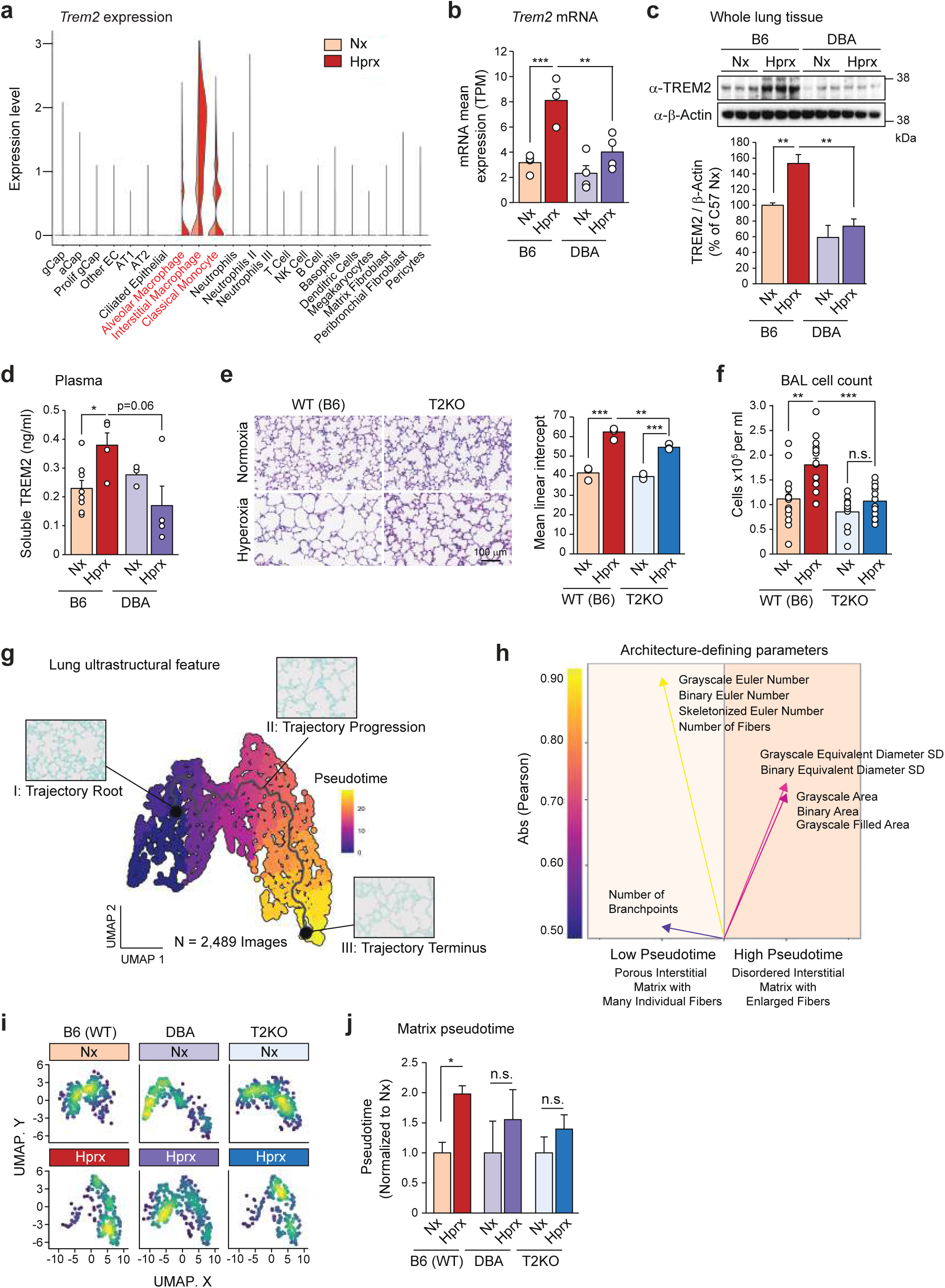
*Trem2* deletion in myeloid cells protects the developing lung from (neonatal) hyperoxia-induced injury. (a) Single-cell RNA-seq analysis of whole lung tissues from B6 mice exposed to 95% oxygen from P0 to P5 ^1^. (b) *Trem2* mRNA expression in the lungs of B6 and DBA mice. Bar graphs show mean ± SEM (n = 3-4 for each group). ANOVA was performed followed by Tukey’s post hoc comparison. \*\**p*-adj < 0.01 and \*\*\**p*-adj < 0.001. (c) Immunoblots for TREM2 protein in whole lung tissues collected on P14 from B6 and DBA mice. Bar graphs show mean ± SEM (n = 3 for each group). ANOVA was performed followed by Tukey’s post hoc comparison. \*\**p* < 0.01. Uncropped images of the blots are shown in Extended Data Fig. 7. (d) Soluble TREM2 levels in the plasma of B6 and DBA mice on P14 after neonatal hyperoxia exposure. Bar graphs show mean ± SEM (n = 3-8 per group). ANOVA was performed followed by Tukey’s post hoc comparison. \**p* < 0.05. (e) H&E staining was performed to assess the alveolar complexity at P14 in WT (B6) mice (left panels) and T2KO mice (right panels) with scale bars denoting 100 μm. The bar graph represents the results of the quantification of alveolar simplification using mean linear intercept (same WT (B6) samples as Fig. 1b). Data are mean ± SEM (n = 3-4 for each group). ANOVA was performed followed by Tukey’s post hoc comparison. \*\**p* < 0.01 and \*\*\**p* < 0.001. (f) Bronchoalveolar lavage (BAL) was performed at P14 in WT (B6) and T2KO hyperoxia-exposed and normoxic control mice (n = 14 for WT (B6) and n = 11-13 for T2KO mice). ANOVA was performed followed by Tukey’s post hoc comparison. \*\**p* < 0.01 and \*\*\**p* < 0.001. n.s.: not significant. (g) Matrix ultrastructural analysis of lung tissues of neonatal hyperoxia-exposed B6 (WT), DBA and T2KO mice on P14. Manifold of hyperoxia-induced pathological architecture with higher pseudotime representing increasingly disrupted architecture and alveolar simplification. Tissue images show representative tiles along the pseudotime trajectory. (h) Architecture-defining parameters. Identification of top 5 individual matrix features associated with low pseudotime (normal-like interstitium) and high pseudotime (more aberrant interstitium), based on Pearson correlations. Parameters are displayed as the absolute value of the Pearson coefficient (i.e. magnitude of correlation, as shown by height of arrows on y-axis), with all *p* < 0.001. (i) Visualization of hyperoxia-exposed and normoxic control lung tissues of B6 (WT), DBA and T2KO mice. Hyperoxia-exposed lungs from DBA and T2KO mice with less lung matrix remodeling localize near the root point of the pseudotime trajectory. (j) Bar graphs of matrix pseudotime for hyperoxia-exposed and normoxic control lung tissues of B6 (WT), DBA and T2KO mice. The difference in pseudotime normalized to normoxic control lungs is shown. Data are mean ± SEM (n = 3 for each group). ANOVA was performed followed by Tukey’s post hoc comparison. \**p* < 0.05. n.s.: not significant. Each dot represents one mouse (b, d, e, f). See also Extended Data Fig. 2.

To test if TREM2 expression is differentially modulated in the hyperoxia-sensitive B6 and resistant DBA mice, we measured gene and protein expression in the whole lung. *Trem2* mRNA was only upregulated in the lungs of hyperoxia-sensitive B6 mice but not in the resistant DBA mice (Fig. 2b). Similarly, we found a significant increase in TREM2 protein levels in the lungs of hyperoxia-exposed B6 mice, while TREM2 protein levels did not change significantly in the lungs of DBA mice (Fig. 2c). Since TREM2 can be cleaved from the cell surface and released in the blood, we measured soluble TREM2 levels in the plasma. While we detected a significant increase in soluble TREM2 levels in the plasma of B6 mice, the levels in DBA mice remained unchanged (Fig. 2d). Of note, the coding sequence of *Trem2* gene is identical in B6 and DBA mice and baseline *Trem2* gene and its protein expression is comparable in the two strains of mice (Fig. 2b, c). There are 5 non-coding SNPs identified in the *Trem2* gene between the two strains that are of uncertain significance with respect to their potential effects on *Trem2* regulation (Extended Data Fig. 2e).

To determine the functional importance of TREM2 in lung myeloid cells during neonatal hyperoxia-induced injury, we exposed *Trem2*-deficient (T2KO) mice and wild-type (WT) controls in the B6 background to hyperoxia in the neonatal period and analyzed the severity of lung injury. Remarkably, T2KO mice preserved a developmentally appropriate lung architecture similar to that of DBA mice (Fig. 1b), while the WT (B6) controls developed marked alveolar simplification (Fig. 2e). To quantify the observed differences in lung structure, we measured mean linear intercept in these mice and found a significant decrease in mean linear intercept in neonatal hyperoxia-exposed T2KO mice compared to WT controls (Fig. 2e). Differences in lung morphology were not altered by differences in growth as both WT and T2KO mice lost similar body weight during hyperoxia exposure (Extended Data Fig. 2f). This is important because caloric intake can modify lung development.

To further characterize the phenotypic difference between T2KO and WT mice, we performed BAL in the hyperoxia-exposed pups and normoxic littermate controls. Hyperoxia exposure resulted in increased cellularity of the BAL fluid in WT mice but not T2KO mice (Fig. 2f). These data suggest that TREM2 expression in myeloid cells is not merely a hyperoxia-associated marker but an essential driver of the injury and abnormal lung development. In the absence of TREM2 signaling in myeloid cells, the hyperoxia-elicited immune response was modified in a beneficial way to support normal lung development.

To further investigate the differences in hyperoxia-induced injury in B6 (WT), DBA, and T2KO mice, we performed an ultrastructural analysis of the lung interstitium to quantify extracellular matrix features using an algorithm developed to analyze fibrotic tissue ^26, 27^. We used picrosirius red to stain collagen fibers and analyzed a total of 2489 images obtained across the three mouse strains (WT (B6), T2KO, DBA) and two experimental conditions (Nx, Hprx). From these images we measured 147 collagen fiber features, including length, width, alignment, and porosity to capture the architecture of the lung interstitium. The obtained high-dimensional fiber feature matrix was reduced by uniform manifold approximation and projection (UMAP) to visualize differences in overall lung interstitial architecture (Fig. 2g). Furthermore, a trajectory that connects datapoints based on similarity in ultrastructural parameters was computed and pseudotime scores were assigned based on relative deviation from a root point that was defined as baseline using normoxic lungs. Pseudotime analysis resulted in three general patterns of lung architecture with the first one representing normal lung (low pseudotime). Increasing pseudotime was associated with increasingly disrupted lung interstitium and alveolar simplification. The normal lungs were associated with low pseudotime values (Fig. 2h, left panel). On the other hand, high pseudotime was defined by disordered, geometrically variable, and enlarged extracellular matrix fibers that are characteristic of interstitial fibrosis (Fig. 2h, right panel). We found marked differences in the ultrastructural changes caused by neonatal hyperoxia in the three mouse strains. While the lungs of hyperoxia-exposed B6 mice were characterized by geometrically disordered and enlarged fibers (high pseudotime), T2KO and DBA mice were enriched in fiber features like normal lungs (low pseudotime) (Fig. 2i, j). We did not observe significant differences in lung architecture at baseline between the 3 mouse strains. The pseudotime normalized to normoxic control lungs was significantly increased in B6 mice but not DBA nor T2KO mice (Fig. 2j).

These findings indicate that neonatal hyperoxia exposure induces a spectrum of morphological changes in the developing lung, including alveolar simplification, decreased alveolar number and size, and changes in interstitial architecture representative of fibrosis. Importantly, the degree of lung injury is associated with the inflammatory response and loss of TREM2 signaling in myeloid cells can prevent these structural alterations.

### Trem2 deficiency remodels the transcriptional landscape in the lung to support normal alveolar growth

To investigate mechanisms underlying the resilience against neonatal hyperoxia-induced injury in T2KO mice, we performed RNA-seq to define the lung transcriptome of WT and T2KO mice under control conditions and in response to hyperoxia exposure. Gene expression profiling of the developing lung at PND14 identified 756 differentially expressed genes (p-adj < 0.05, fold-change > 2) between WT and T2KO mice (Fig. 3a). We identified 90 of these genes were induced by hyperoxia and expressed at significantly higher levels in WT compared to T2KO mice (Fig. 3b, Venn diagram). Gene ontology analysis highlighted enrichment of p53 signaling, autophagy, and cell death related processes in this group of genes (Fig. 3c). Examples of individual genes belonging to the p53 pathway are shown in Fig. 3d. *Bcl2l1*, *Perp*, *Sesn1*, *Trp53inp1*, *Cdkn1a*, and *Fas* were all significantly upregulated by hyperoxia in WT mice but not T2KO mice similar to DBA mice (Fig. 1g). To verify these results at the protein level, we measured cleaved caspase 3 in cytosolic fractions from whole lungs and found a significantly higher level of induction in WT mice compared to T2KO mice (Fig. 3e). In addition to exhibiting reduced expression of p53 pathway genes, T2KO mice upregulated cell proliferation-related genes such as *Ccnd1*, *Vegfb*, *Wnt3a*, and *Sfrp2* (Fig. 3f). Several inflammation-related genes showed a similar pattern of regulation as the p53 pathway genes; they were upregulated in WT mice but remained unchanged in T2KO mice (Fig. 3g). For example, cytokine receptors encoding genes *Il6ra* and *Il20b* were increased in WT mice. IL6 is a pleiotropic cytokine known to regulate cell growth and immune responses in the lung. In addition to inflammation promoting genes, genes encoding CD73 and CD39, whose primary function is to convert extracellular ATP to adenosine and hereby limit an excessive immune response in various pathophysiological events including lung injury, were elevated in WT mice. Taken together, these results show that *Trem2* deficiency finetunes the innate immune response and prevents the activation of the p53 pathway in the neonatal lung after hyperoxia exposure and hereby preserves normal alveolar development. Of note, the transcriptomic changes observed in T2KO mice recapitulate the ones observed in the hyperoxia resistant DBA mice (Extended Data Fig. 3a) and exhibit less inflammation and less cell death and apoptosis (Fig. 1g, Extended Data Fig. 3b).

**Fig. 3.**
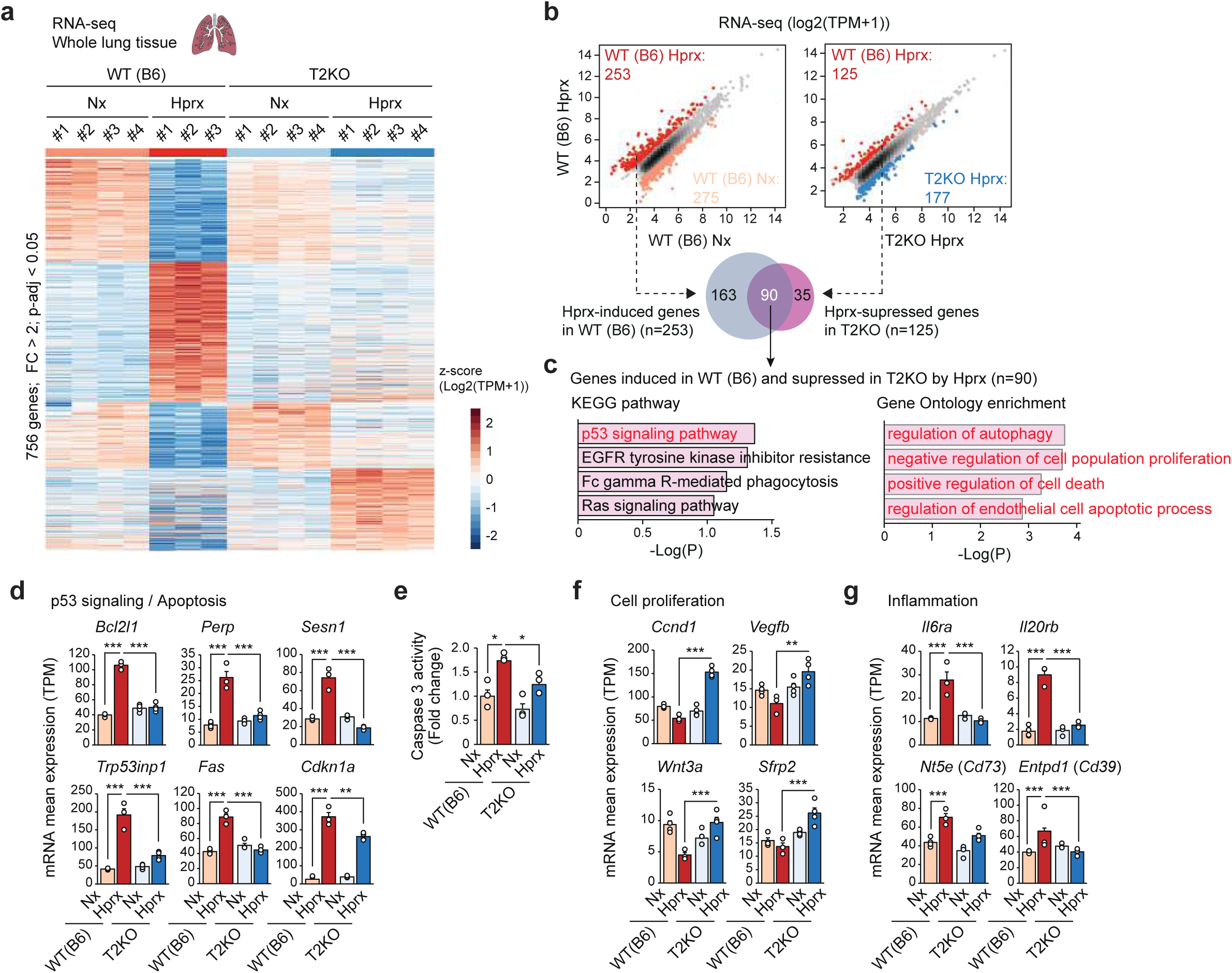
Apoptosis and cell proliferation related gene programs are differentially activated in the lungs of WT (B6) and *Trem2* deficient (T2KO) mice. (a) RNA-seq in whole lung tissues of WT (B6) (same samples as Fig. 1d) and T2KO mice at P14. Heatmap shows the 756 differentially expressed genes (FC > 2; *p*-adj < 0.05) comparing hyperoxia-exposed WT and T2KO mice with the normoxic controls. (b) Scatterplots of RNA-seq data in whole lung tissues showing hyperoxia-regulated gene expression in WT (B6) mice (left panel, same as Fig. 1e) and T2KO mice-regulated gene expression in the hyperoxic conditions (right panel). Dark red dots in left panel: significant hyperoxia-induced genes in WT mice, light red dots in left panel: significant hyperoxia-suppressed genes in WT mice, dark red dots in right panel: significant T2KO-suppressed genes in the hyperoxic conditions, dark blue dots in right panel: significant T2KO-induced genes in the hyperoxic conditions (FC > 2; FDR < 0.05). Venn diagram shows the overlap between hyperoxia-induced genes in WT mice (n = 253) and hyperoxia-suppressed genes in T2KO mice (n = 125). (c) KEGG pathway (left panel) and Gene Ontology enrichment (right panel) analysis of the 90 genes that were induced in WT and suppressed in T2KO by hyperoxia. (d) Bar plots for expression of representative genes belonging to the p53 signaling pathway. Data are mean ± SEM. DSeq2 was used for comparisons. \*\**p*-adj < 0.01 and \*\*\**p*-adj < 0.001. (e) Caspase 3 activity measured in cytosolic fractions from whole lung tissues from B6 (same as Fig. 1h) and T2KO mice harvested on P14. Data are mean ± SEM (n = 3 for each group). ANOVA was performed followed by Tukey’s post hoc comparison. \**p* < 0.05. (f) Bar plots for expression of representative genes belonging to the cell proliferation pathway. Data are mean ± SEM. DSeq2 was used for comparisons. \*\**p*-adj < 0.01 and \*\*\**p*-adj < 0.001. (g) Bar plots for expression of representative inflammatory genes. Data are mean ± SEM. DSeq2 was used for comparisons. \*\*\**p*-adj < 0.001. Each dot represents one mouse (d, e, f, g). See also Extended Data Fig. 3.

### Hyperoxia directly affects gene expression in myeloid cells

Hyperoxia, known to induce lung inflammation, can activate myeloid cells through various mechanisms ^28, 29, 30^. This can occur indirectly, by damaging lung epithelial and endothelial cells eliciting a myeloid response, or by directly activating the myeloid cells. To investigate the direct effects of hyperoxia on myeloid cells, we isolated bone marrow cells from B6, DBA, and T2KO mice, differentiating the cells into bone marrow-derived macrophages (BMDMs) using established protocols (Fig. 4a) ^31^. We exposed half of the BMDM cultures to hyperoxia (95% oxygen) for 24 hours, while maintaining the other half in normoxic conditions (21% oxygen) as controls. The 24-hour hyperoxia exposure period was chosen based on a preliminary time-course experiments to achieve sufficient activation of TREM2 (Extended Data Fig. 4a).

**Fig. 4.**
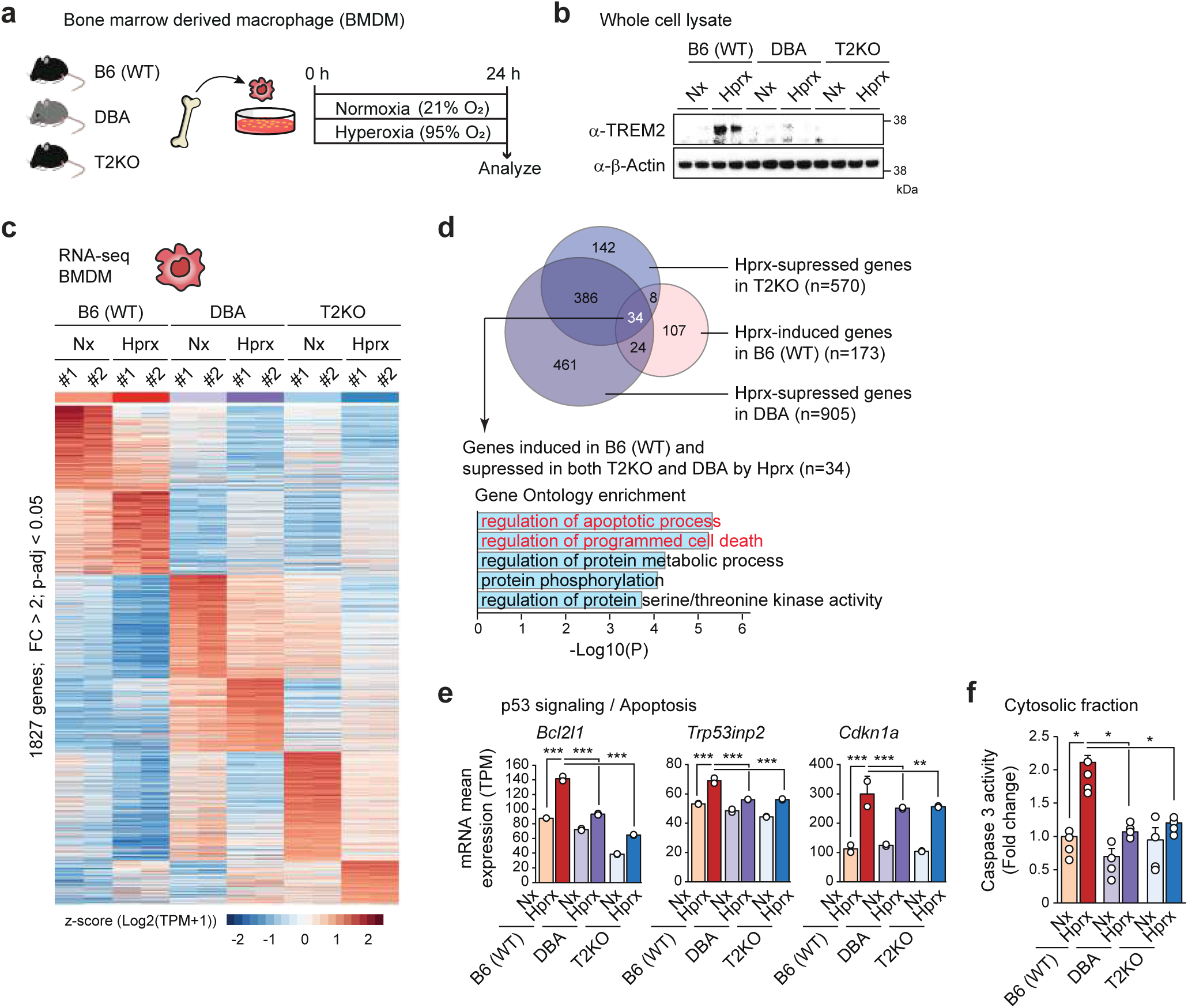
*In vitro* hyperoxia exposure (95% oxygen) of bone marrow-derived macrophages (BMDMs) increases TREM2 expression. (a) BMDMs were obtained from B6 (WT), DBA and T2KO mice and exposed to hyperoxia (95%) for 24 h. (b) Immunoblots of TREM2 protein in whole cell lysates of hyperoxia-exposed BMDMs obtained from B6 (WT), DBA and T2KO mice and the normoxic controls. Uncropped images of the blots are shown in Extended Data Fig. 7. (c) RNA-seq of hyperoxia-exposed BMDMs from B6 (WT), DBA and T2KO mice and the normoxic controls. (d) Venn diagram showing the overlap between hyperoxia-induced genes in B6 (WT) BMDMs (n = 173) and hyperoxia-suppressed genes in T2KO (n = 570) and DBA (n = 905) BMDMs. Gene Ontology analysis of 34 genes induced by hyperoxia in B6 (WT) BMDMs and suppressed in T2KO and DBA BMDMs. (e) Bar plots for expression of representative genes belonging to the p53 signaling pathway. Data are mean ± SEM. DSeq2 was used for comparisons. \*\**p*-adj < 0.01 and \*\*\**p*-adj < 0.001. (f) Caspase 3 activity measured in cytosolic fractions from BMDMs from B6 (WT), DBA and T2KO mice. Data are mean ± SEM. (n = 4 for each group). ANOVA was performed followed by Tukey’s post hoc comparison. * *p* < 0.05. Each dot represents one mouse (e, f). See also Extended Data Fig. 4.

Our initial analysis of TREM2 expression revealed that hyperoxia induced TREM2 protein levels in BMDMs from hyperoxia-sensitive B6 mice, but not in those from hyperoxia-resistant DBA mice. T2KO BMDMs were included as negative controls (Fig. 4b). To elucidate the underlying mechanisms and strain-specific differences, we performed RNA-seq on BMDMs from all three mouse strains. There were significant strain-specific transcriptomic differences, with 1827 genes differentially expressed among the strains under hyperoxia (Fig. 4c).

To identify protective gene programs, we focused on 34 genes that were upregulated by hyperoxia in the sensitive B6 strain but downregulated in the resistant DBA and T2KO mice relative to WT (Fig. 4d). Gene ontology analysis indicated an enrichment for genes involved in apoptotic processes and programmed cell death within this group. Consistent with whole lung transcriptomic changes, genes such as *Bcl2l1*, *Trp53inp2*, and *Cdkn1a* were significantly more induced in BMDMs from B6 mice under hyperoxia compared to DBA and T2KO mice (Fig. 4e). To confirm these transcriptomic findings at the protein level, we assessed cleaved caspase 3 activity in BMDM cytosolic fractions from all three strains. Hyperoxia exposure increased cleaved caspase 3 activity in BMDMs from B6 mice, but not in those from DBA or T2KO mice (Fig. 4f).

### Neonatal hyperoxia alters p53 binding landscape in B6 mice but not in the resistant DBA and T2KO mice

Analysis of the RNA-seq data indicated that the p53 pathway was a significant determinant of lung injury. Since p53 target gene regulation is dependent on its role as a transcription factor, we measured p53 protein levels in the nuclear fraction of whole lungs. We show a significant increase of p53 protein in the nucleus of the whole lung from hyperoxia-sensitive B6 mice as compared to hyperoxia-resistant DBA (Fig. 5a, left panel) and T2KO (Fig. 5a, right panel) mice. To better understand the role of p53 in neonatal hyperoxia-induced lung injury, we employed chromatin immunoprecipitation followed by sequencing (ChIP-seq) to map the genome-wide binding sites of p53 and elucidate its downstream target genes. We performed replicate ChIP-seq analysis in whole lungs harvested from both hyperoxia-exposed and control neonatal B6, DBA and T2KO mice (Extended Table 1) and used the Irreproducible Discovery Rate method to define confident peaks. Hundreds of p53 peaks were identified in all three strains of mice (Extended Data Fig. 5a). To identify p53 binding sites that are hyperoxia-dependent, differential binding analysis comparing hyperoxia-exposed and control samples detected 53 hyperoxia-induced loci in B6 mice (p-adj < 0.05, fold-change > 2). Hyperoxia-induced changes in p53 were much more modest in the hyperoxia-resistant DBA and T2KO mice (Fig. 5b). Further analyzing the hyperoxia-induced p53 cistrome in B6 mice, motif enrichment analysis revealed that the most strongly enriched binding motif was the canonical p53 motif (p53, p63, and p73 recognize what is essentially the same motif) (Fig. 5c). In addition, the motifs recognized by zinc finger 416 (ZF416), a transcription factor reported to negatively regulate cell growth and zinc finger protein 809 (ZFP809), a transcription factor required to establish epigenetic silencing of ancient endogenous retrovirus (ERVs) during embryonic development ^32^, were enriched at p53 binding sites. Interestingly, while p53 binding sites were found to be mainly in introns and intergenic regions, 15% of binding sites were in long terminal repeats (LTRs), DNA sequences that form a retrotransposon, an endogenous retrovirus, or a retroviral provirus (Extended Data Fig. 5b). Furthermore, we found that p53 binding sites were enriched in TEAD motifs, which bind a transcription factor that plays a critical role in the Hippo pathway and is known to restrict proliferation and to promote apoptosis ^33^.

**Fig. 5.**
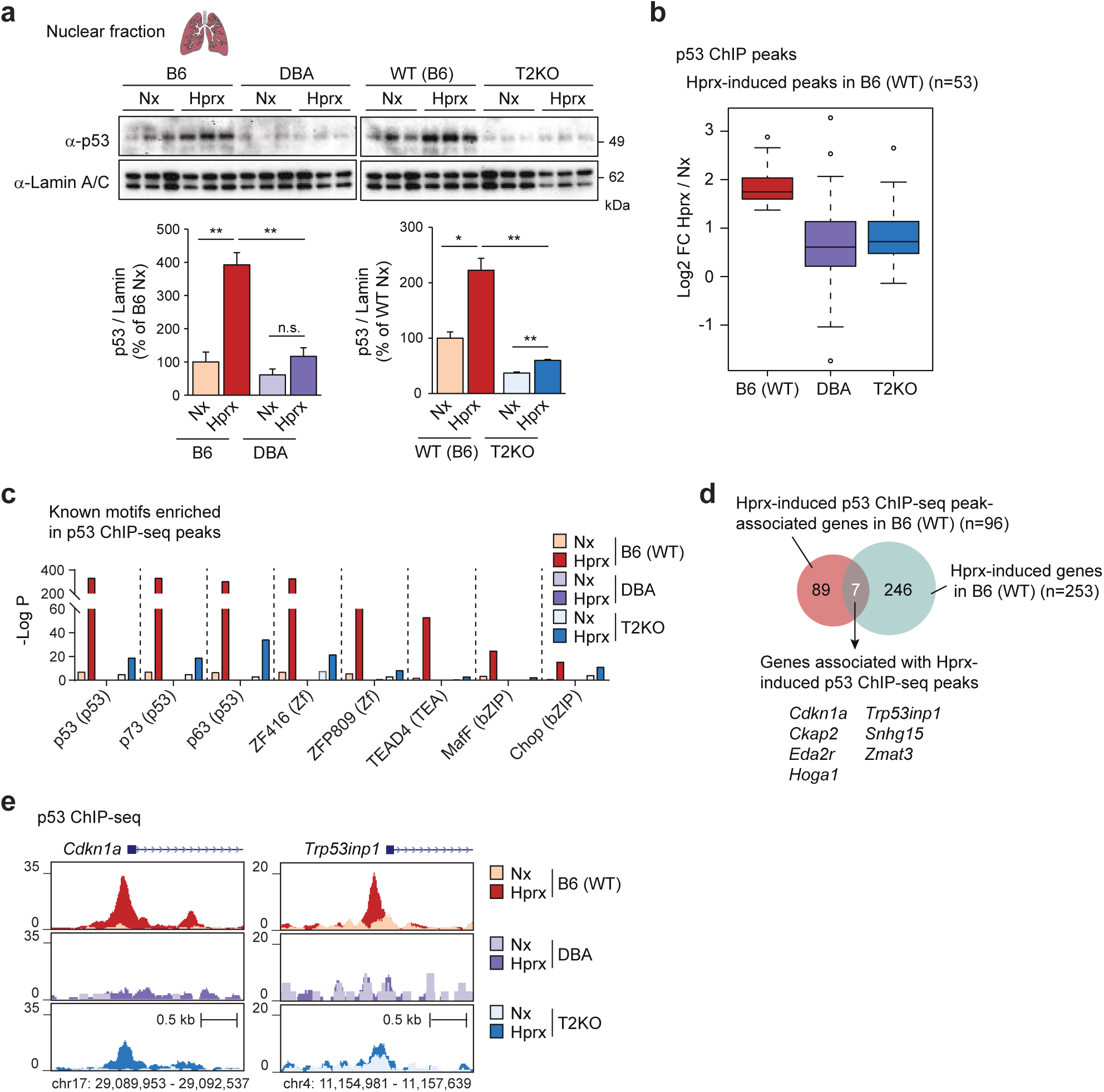
Divergent p53 protein expression and DNA binding pattern in WT (B6), DBA and T2KO mice. (a) Immunoblots for p53 protein in the nuclear fraction from whole lung tissues of neonatal hyperoxia-exposed and control B6 and DBA mice (left panel) and WT (B6) and T2KO mice (right panel) harvested on P14. Data are mean ± SEM (n = 3 for each group). ANOVA was performed followed by Tukey’s post hoc comparison. \**p* < 0.05 and \*\**p* < 0.01. n.s.: not significant. (b) Boxplot for p53 ChIP-seq peaks in whole lung tissues from B6 (WT), DBA and T2KO mice. The 53 hyperoxia-induced peaks (FC > 2; *p* < 0.05) in lungs of B6 (WT) mice were used for the analyses for DBA and T2KO mice. (c) Known motifs enriched in p53 ChIP-seq peaks in whole lung tissues using a GC-matched genomic background. (d) Venn diagram showing the intersection of hypoxia-induced genes in whole lung tissues of B6 (WT) mice (n = 253, FC < 2; FDR < 0.05, Fig. 1e) and genes found near p53 ChIP-seq peaks (n = 96, 25 kbp from TSS) in whole lung tissues of B6 (WT) mice. *p* = 2.4E-8 (Cumulative binomial distribution). (e) Genome browser track showing p53 ChIP-seq peaks in the vicinity of the *Cdkn1a* and *Trp53inp1* genes. See also Extended Data Fig. 5.

To link p53 binding to gene expression changes, we performed an integrative analysis of ChIP-seq and RNA-seq data. This analysis showed that hyperoxia-induced genes are 10 times more likely to have a p53 binding site (Fig. 5d). Intersecting the 253 genes induced by hyperoxia in B6 mouse lungs (Fig. 1e) with the genes bound by p53 ChIP-seq peaks (< 25 KB from TSS), we identified 7 genes (Fig. 5d). The group of hyperoxia-induced genes directly bound by p53 includes *Cdkn1a* and *Trp53inp1*. The genome browser tracks for these two genes are shown in Fig. 5e (https://genome.ucsc.edu/s/Cbenner/Sajti%2D240815%2DReviewerTracks). While only a small number of hyperoxia-regulated genes were directly bound by p53, this association was highly specific as hyperoxia-induced genes are 10 times more likely to have a p53 binding site near their promoters than non-regulated genes.

Taken together, ChIP-seq analysis of p53 in the lung highlights enrichment in apoptotic pathway, suggesting a multifaceted role of p53 in orchestrating the cellular response to hyperoxia-induced lung injury. Comparative analysis between different mouse strains (e.g., hyperoxia-sensitive B6 vs. resistant DBA and T2KO) revealed that hyperoxia only altered p53 genomic localization in the hyperoxia-sensitive B6 mice.

### Gene regulation by p53 is similar in whole lung and myeloid cells

The sensitivity of ChIP-seq of whole lungs for p53 is likely to be limited by the extent to which the p53 program is activated among the different cell types of the lung and to be strongly biased by the p53 binding profile in the most abundant responsive cell types, which may not be myeloid cells. This is supported by the lack of macrophage lineage determining factors in the enriched motifs. To elucidate the role of TREM2 signaling in orchestrating a protective innate immune response in neonatal hyperoxia-induced lung injury, we analyzed the cell-specific role of p53 signaling in myeloid cells. First, we assessed p53 translocation to the nucleus in BMDMs following hyperoxia exposure. We observed a significant increase in nuclear p53 protein levels in hyperoxia-exposed BMDMs from B6 mice. In contrast, genetic deletion of TREM2 did not affect p53 protein levels in BMDM nuclei after hyperoxia exposure (Fig. 6a).

**Fig. 6.**
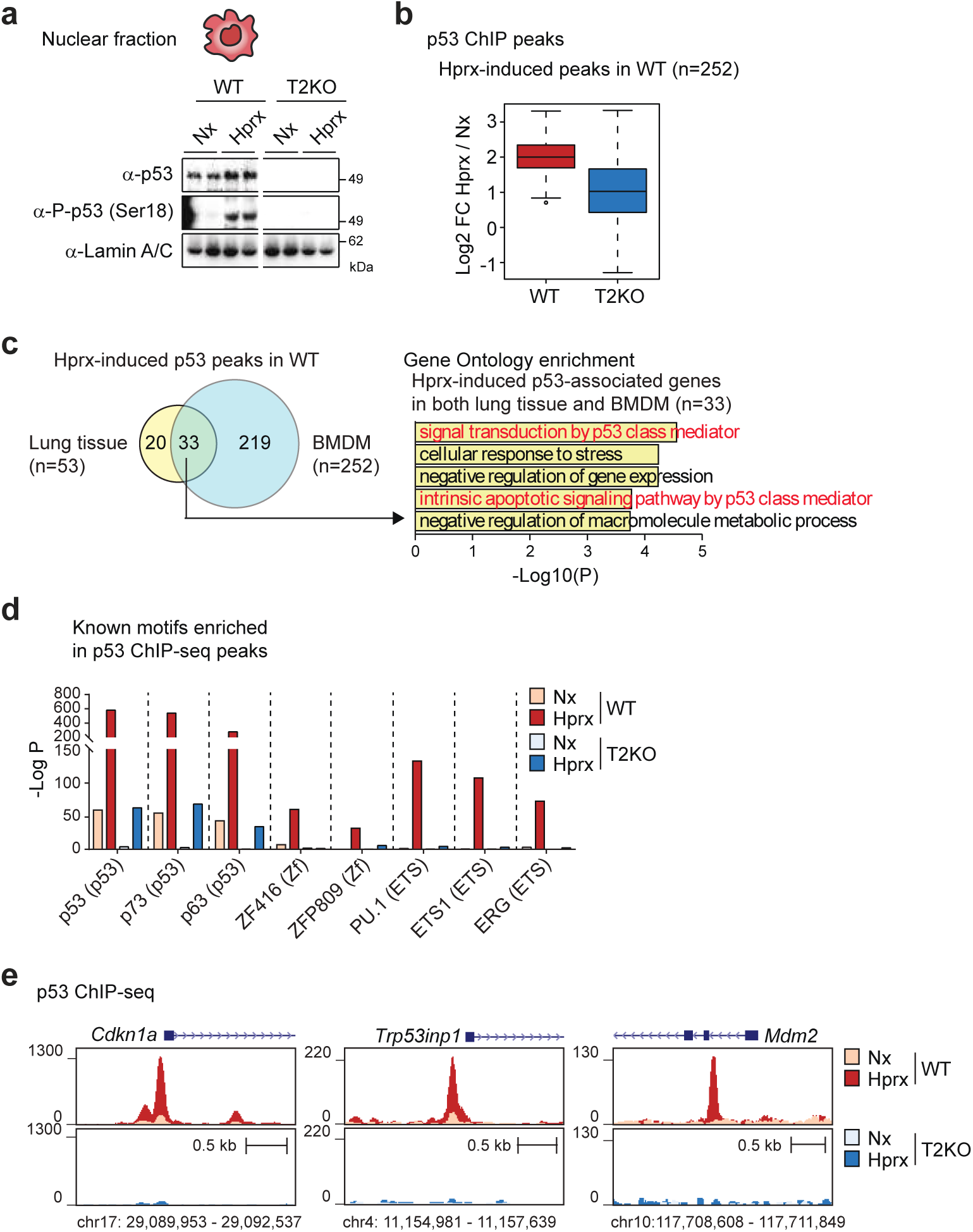
Divergent p53 DNA binding pattern in BMDMs of B6 and T2KO mice. (a) Immunoblots of total p53, phospho-p53 (Ser18), and Lamin A/C in the nuclear fraction of hyperoxia-exposed (24 h) BMDMs from WT and T2KO mice. Uncropped images of the blots are shown in Extended Data Fig. 7. (b) Boxplot for p53 ChIP-seq peaks in BMDMs from WT and T2KO mice. The 252 hyperoxia-induced peaks (FC > 2; *p* < 0.05) in BMDMs of WT mice were used for the analyses for T2KO mice. (c) Venn diagram showing the overlap of hyperoxia-induced p53 ChIP-seq peaks in the lung tissues and BMDMs of WT mice. (d) Known motifs enriched in p53 ChIP-seq peaks in BMDMs using a GC-matched genomic background. (e) Genome browser track showing p53 ChIP-seq peaks in the vicinity of the *Cdkn1a, Trp53inp1* and *Mdm2* genes. See also Extended Data Fig. 6.

Given that activated p53 coordinates a flexible gene expression program that can lead to cell death or cell-cycle arrest and DNA repair ^34^, we investigated the mechanisms driving divergent outcomes in WT and TREM2-deficient myeloid cells as shown in Fig. 4. It is well-established that post-translational modifications of p53, including multisite phosphorylation, regulate p53-mediated cell fate decisions ^35, 36^. Therefore, we measured the expression of the p53 Ser18 phospho-isomer corresponding to Ser 15 of human p53 (Extended Data Fig. 6a) in the nuclear fraction of mouse BMDMs. It was previously reported that phosphorylation at Ser18 was required for a full p53-mediated response to DNA damage-inducing stimuli ^37^. A significant increase in the p53 Ser18 phospho-isomer was observed in BMDMs from B6 mice after hyperoxia exposure, but not in T2KO mice (Fig. 6a).

To delineate the p53 binding landscape in BMDMs, we performed ChIP-seq analysis (Extended Data Table 1). Thousands of p53 binding sites were identified in BMDMs from both B6 (WT) and T2KO mice (Extended Data Fig. 6b). The relative distribution of hyperoxia-induced p53 ChIP-seq peaks across various genomic elements is shown in Extended Data Fig. 6c. Differential binding analysis revealed 252 hyperoxia-induced binding sites (p-adj < 0.05, fold-change > 2). Notably, hyperoxia had a reduced impact on p53 binding in the absence of TREM2 signaling (Fig. 6b). We observed significant overlap between hyperoxia-induced p53 peaks in whole lungs and BMDMs from B6 mice, with 33 commonly induced p53 peaks associated with genes involved in apoptosis and cellular stress responses (Fig. 6c). Motif analysis identified the canonical p53 motif (p53, p63, and p73 motifs are highly similar) as the most enriched binding motif (Fig. 6d), consistent with findings from the whole lungs (Fig. 5c). Myeloid cell-specific p53 motifs included PU.1, ETS, and ERG (Fig. 6d). Genomic distribution of p53 binding sites in BMDMs and whole lung was similar with most of the p53 binding sites in introns and intergenic regions (Extended Data Fig. 6c). The group of hyperoxia-induced genes directly bound by p53 includes *Cdkn1a*, *Trp53inp1*, and *Mdm2* (Extended Data Fig. 6d). The genome browser tracks for these three genes are shown in Fig. 6e (https://genome.ucsc.edu/s/Cbenner/Sajti%2D240815%2DReviewerTracks). These results indicate that hyperoxia regulation of p53 in BMDMs is cell autonomous, or at the very least, it indicates that TREM2 has a direct interaction with the p53 pathway.

## Discussion

BPD remains a profound challenge in neonatal care despite significant advancements in respiratory support for premature infants. This condition not only affects immediate neonatal health but also predisposes individuals to long-term respiratory complications. Intriguingly, there exists considerable variability in BPD severity among premature infants who receive similar standards of care, suggesting that factors beyond clinical management, such as genetic predispositions and molecular pathways, play a critical role in disease progression.

Our study leverages genetically diverse mouse models to explore the underlying mechanisms contributing to varying susceptibilities to hyperoxia-induced lung injury, a common factor in the development of BPD. The findings underscore the activation of the immune system and the p53 pathway as pivotal contributors to severe disease phenotypes. Notably, our research highlights the involvement of TREM2 signaling in myeloid cells during hyperoxic conditions as a key driver of the inflammatory response that leads to detrimental outcomes such as alveolar simplification and fibrosis.

TREM2 is a transmembrane receptor selectively expressed on the surface of myeloid cells, including macrophages and microglia, and has been extensively studied in the context of neurodegenerative diseases. However, our findings extend its significance to pulmonary health, revealing a novel role in lung pathology. During hyperoxia exposure, TREM2 signaling exacerbates the inflammatory response in the neonatal lung, which impairs normal alveolar development and promotes fibrosis. The remarkable protective effect observed with the genetic deletion of Trem2, which resulted in almost complete prevention of lung tissue injury, underscores the potential of TREM2 as a therapeutic target for mitigating BPD severity.

In addition to its membrane bound form, TREM2 also has a soluble form that can be detected in blood ^18, 38^. We show that soluble TREM2 (sTREM2) was increased in circulating blood of hyperoxia sensitive B6 mice but not in the hyperoxia-resistant DBA mice. The functional difference between the membrane bound and soluble form is a topic of ongoing investigations, sTREM2 might be a potential biomarker for disease severity in BPD.

Previous studies investigating TREM2 in lung disease have shown a disease promoting role for both the membrane bound and soluble forms in murine pneumonia ^20, 21^. In humans, increased sTREM2 was detected in the bronchoalveolar lavage fluid of patients with asthma and sarcoidosis as compared to healthy controls ^22, 23^. Taken together, these studies suggest a role for TREM2 in a variety of lung diseases. The disease-promoting role of TREM2 described in lung disease contrasts with its protective role in Alzheimer’s disease, underscoring the importance of studying TREM2 effects in a tissue and disease specific way.

Macrophages are central to both the resolution of organ injury and the pathogenesis of pulmonary fibrosis. They orchestrate the immune response and produce reactive oxygen species, contributing to tissue damage and fibrosis when dysregulated ^39, 40^. Our study implicates TREM2 as a critical modulator of macrophage activity in the lungs under hyperoxic stress, linking it to the initiation and perpetuation of an injurious inflammatory response. This aligns with the broader understanding of macrophage functions in organ injury and repair, where their role can be both protective and destructive depending on the context and regulatory mechanisms involved.

TREM2 is known to interact with a variety of ligands, which are typically associated with damaged or stressed cells and can trigger immune responses. Some of the known ligands for TREM2 include lipids such as phosphatidylserine and phosphatidylcholine, which are components of cell membranes and can become exposed on the surface of apoptotic or necrotic cells ^41, 42^. While it is well established that hyperoxia exposure results in epithelial and endothelial cell death in the lung, further studies are needed to elucidate the precise ligands that trigger the TREM2 pathway in myeloid cells.

BPD has a strong genetic background and based on twin studies heritability is estimated to be between 50-80% ^43^. However, it has been challenging to pinpoint the genes that drive the disease. GWAS studies published to date include relatively small sample sizes and a heterogeneous group of patients. Nevertheless, a consistent finding across these studies is that BPD predisposing genes are related to the inflammatory response. These genes included *CRP*, *NUAK1*, *SELL*, *VNN2*, and *CHST3*, four of them related to cell adhesion ^44^. Twin studies have demonstrated a clear genetic component for BPD ^45, 46^. However, studies examining single nucleotide polymorphisms (SNPs) genome wide or in candidate genes were unable to pinpoint any specific gene that would be the cause of BPD ^47, 48^. There is evidence that SNPs in candidate genes affecting the immune response and oxidative stress are significant predisposing factors for BPD ^44^. For example, SNPs in the C-reactive protein, a well-known inflammatory molecule was associated with BPD ^49^. Sequence variants of the antioxidant response gene *NQO1* (encoding NAD(P)H quinone dehydrogenase1) were associated with increased incidence of BPD in two independent studies ^50, 51^. The negative findings of genome wide association studies can be explained, at least partially, by the small sample sizes. Second, many disease susceptibility genes are cell-type specific and gene expression in aggregate peripheral blood cells might not capture disease relevant gene regulatory networks in the affected organ. Third, most of the natural genetic variation associated with disease risk resides in non-coding, regulatory regions of the genome ^52^.

More recently, using single-cell RNA sequencing, we and others have shown that neonatal hyperoxia exposure significantly alters not only the composition of the myeloid cell compartment but also the gene expression with a unique signature in each myeloid cell subset ^16, 17, 53^. Lung macrophage and monocyte populations, the first responders to injury were among the cells with the largest transcriptomic changes. Importantly, the cell-type specific response of myeloid cells to hyperoxia-induced injury occurs during a dynamic period of lung development. During the saccular and alveolar phases of lung development not only the structural cells (e.g. epithelial, mesenchymal cell) but also the myeloid cells are undergoing important developmental changes ^54^. Myeloid cells can interact with virtually every cell in the lung, and analysis of the complex cellular interactions in hyperoxia-sensitive mice suggests that inflammatory signaling is a main driver of the injury ^16, 17^. However, an innate immune response, when properly executed is crucial for the healing process. Therefore, a successful therapy can not entirely suppress the immune response. Here we show, that modulating the immune response by eliminating TREM2 signaling in myeloid cells supports a regenerative response and prevents a dysregulated one that enhances injury.

The p53 pathway is induced in cell types in the lung that do not express TREM2. This implies that the TREM2-dependent activation of p53 in myeloid cells leads to a TREM2-independent mechanism of p53 activation in non-myeloid cells that is associated with inflammation and tissue pathology. Determining the mechanisms by which the TREM2-p53 signaling axis in myeloid cells results in these effects in other structural lung cell types will be an important objective. While it is known that p53 is induced in the hyperoxia exposed lung ^55, 56^, it’s cell type-specific role is not fully elucidated. P53 is best-known for its role in tumor suppression ^57, 58^. However, new evidence shows that inappropriate activation of p53 during development can trigger a wide range of defects involving multiple organs including the lung ^59^. Importantly, timing and degree of p53 activation can have a substantial impact on phenotypic outcomes ^60^. In addition, being able to sense a gamut of stress signals and detect foreign genomes, p53 evolved to become an important part of the innate immune system ^61, 62, 63^. Since the pathophysiology of BPD involves both inflammation and development, relevant mechanisms can only be uncovered by accounting for both processes. The role of p53 in both development and innate immunity makes this an interesting signaling node that potentially integrates stress signals to simultaneously modify inflammatory responses and developmental processes. At a molecular level, p53 can be activated by a large variety of signals including DNA damage and oxidative stress. Upon activation p53 induces a large network of genes, resulting in a highly connected signal transduction complex ^64^. The best-studied pathways induced by p53 are cell cycle arrest followed by DNA repair or apoptosis.

In conclusion, our findings provide novel insights into the molecular pathways influencing BPD severity, particularly the role of TREM2 signaling in myeloid cells. The prevention of lung injury through TREM2 deletion offers a promising avenue for therapeutic intervention, potentially transforming the management and outcomes of premature infants at risk for BPD. Future research should focus on further elucidating the complex interplay between genetic factors, immune responses, and environmental exposures in BPD pathogenesis, paving the way for targeted therapies that can mitigate this debilitating condition.

Our results emphasize the importance of a balanced lung innate immune response to early life insults such as hyperoxia to optimize normal lung development in premature infants and to improve health outcomes later in life.

## Methods

### Animal studies

All animal procedures were in accordance with University of California, San Diego research guidelines for the care and use of laboratory animals. Mice were maintained under a 12 h light/12 h dark cycle at constant temperature (20-23°C) with free access to food and water. Animals were fed a normal chow diet (T8604, Envigo). Trem2 knockout (Strain #: 027197), C57BL/6J (Strain #: 000664), and DBA (Strain #: 000671) mice were purchased from Jackson Laboratory.

### Neonatal hyperoxic lung injury

For the neonatal hyperoxia experiments, we exposed neonatal mice within 24 h of birth to 75% oxygen until postnatal day 14 (P14). The control pups were raised in room air. For each experiment, neonatal mice of the appropriate strain or genotype were randomized to the hyperoxia, or controls groups based on sex to assure an equal distribution of male and female mice in both experimental groups. Litter size was kept at 5-7 mice. To prevent respiratory distress in the dams, they were exchanged between the experimental and control cages every other day. The weight of the pups was measured every other day.

### Lung harvest

Mice were euthanized with CO_2_. After respiratory movements have ceased, mice were transferred to a dissection area, the abdomen and chest were opened through a midline incision. The aorta was cut, and the lungs were perfused with ice cold PBS through the right ventricle until the lungs were cleared of blood. Lungs were gently removed from the thorax, the lobes were dissected from the trachea, placed in sterile tubes and snap frozen in liquid nitrogen. Tissue was stored at −80°C until used for downstream RNA-seq, ChIP-seq analysis, and protein measurements. For histology, after perfusion of the lungs, a knot was tied loosely around the trachea with size 4/0 silk suture and 24G cannula was inserted into the trachea. Through the cannula, the lungs were inflated with 4% paraformaldehyde to a pressure of 20 cm H_2_O. After the lungs equilibrated to this pressure, the cannula was removed while the knot was tightened to prevent loss of pressure. The inflated lungs where dissected from the thorax and placed in 4% paraformaldehyde for overnight fixation at 4°C.

### Histology and ultrastructural analysis

Lungs inflated at a pressure of 20 cm H_2_O and fixed in 4% paraformaldehyde, were embedded in paraffin and sectioned. Hematoxylin and eosin staining was performed to assess tissue morphology, to measure mean linear intercept (MLI) measurements. For the MLI measurements, full slides were scanned with the Olympus VS200 microscope scanner at 20x and MLI was calculated by overlaying a 50 μm grid on top of 8 representative sections from each mouse, spread over the entire tissue area. Intercepts were measured as points where the tissue crossed over the grid overlay. For each representative section in which intercepts were counted the total measured length was divided by the total number of intercepts counted. This final operation gave the MLI for measurement of alveolar size in μm per intercept.

For ultrastructural analysis, paraffin embedded tissue slides were stained with picrosirius red. A total of 2489 picrosirius red-stained histology image tiles were processed using a matrix ultrastructural algorithm previously utilized for fibrotic tissue analysis ^26, 27^. In brief, whole-slide images (WSI) were programmatically tiled to 300×200 µm in *magick* based on prior utilization ^27^, then processed using the MATLAB Image Processing Toolbox (MathWorks Inc.) by red/green/blue (RGB) normalization and color deconvolution as established by Ruifrok et al. ^65, 66^. Matrix staining (green) was further processed by noise reduction using a Wiener filter in 3×3 pixel neighborhoods to smooth low-variance regions and preserve fiber edges, then binarized by *im2bw*, processed using erosion filters, and skeletonized by *bwmorph*. For each image tile, a set of 147 local (e.g. fiber diameter, length) and global (e.g. fiber branching, porosity) features was quantified, and the resulting values were reduced by UMAP and ported to *RStudio* for downstream analysis ^26, 27^. *DDRTree* was applied to fit all tiles to a minimum spanning tree based on stepwise deviations in fiber features, assigning pseudotime scores proportional to each tile’s geodesic distance from the trajectory root ^67, 68^. Pseudotime scores were averaged at the slide-level to determine average ECM architecture, and dataset-level pseudotime was further correlated with individual matrix features using Pearson correlations for post-hoc explanatory analysis.

### Immunoblotting

Whole tissue homogenates or whole cell lysates were prepared as previously described ^69^ with modifications as follows. Pulverized frozen mouse lung tissues or BMDMs were sonicated in cell lysis buffer (50 mM HEPES-KOH (pH 7.9), 150 mM NaCl, 1.5 mM MgCl_2_, 1% NP-40, 1 mM PMSF (Sigma-Aldrich), 1X protease inhibitor cocktail (Sigma-Aldrich) by ultrasound homogenizer (Bioruptor, Diagenode) for 10 min at 4°C. After centrifugation at 15000 g for 10 min at 4°C, the supernatant was used as whole tissue homogenates or whole cell lysates. To isolate the nuclear fraction from frozen mouse lung tissue or BMDMs, the pulverized tissues or BMDMs were homogenized in STM buffer (250 mM Sucrose, 50 mM Tris-HCl (pH 7.4), 5 mM MgCl_2_, 1 mM PMSF, 1X protease inhibitor cocktail) and incubated on ice for 30 min. After centrifugation at 800 g for 15 min at 4°C, the pellet was washed with STM buffer twice and resuspended in NET buffer (20 mM HEPES (pH7.9), 1.5 mM MgCl_2_, 0.5 M NaCl, 0.2 mM EDTA, 20% Glycerol, 1% Triton-X, 1 mM PMSF, 1X protease inhibitor cocktail). The suspension was passed through an 18-gage needle 15 times and sonicated by ultrasound homogenizer for 10 min at 4°C. After centrifugation at 9000 g for 30 min at 4°C, the supernatant was used as the nuclear fraction.

For immunoblotting, aliquots of whole cell lysate or nuclear fraction were boiled at 95°C for 5 min in NuPAGE^TM^ LDS Sample Buffer (Thermo Fisher Scientific) with NuPAGE^TM^ Sample Reducing Agent (Thermo Fisher Scientific), subjected to SDS-PAGE, and transferred to immobilon-P transfer membranes (Merck Millipore). Immunodetection was carried out with the indicated antibodies (Extended Data Table 2) and bound antibodies were visualized with peroxidase-conjugated affinity-purified donkey anti-mouse or anti-rabbit IgG (Dako), or HRP-linked anti-rat IgG (Cell Signaling Technology) using SuperSignalTM West Femto Maximum Sensitivity Substrate (Thermo Fisher Scientific) or Luminate^TM^ Forte Western HRP Substrate (Merck Millipore), and luminescence images were analyzed by ChemiDoc XRS+ System (Bio-Rad Laboratories). Uncropped images of immunoblots are shown in Extended Data Fig. 7.

### Caspase 3 activity assay

Caspase 3 activity in cytosolic extracts from frozen lung tissues was measured using a Caspase-3 Assay Kit (Abcam) according to the manufacturer’s protocol. Briefly, 50 ml of cytosolic extract was mixed with 50 ml of 2X Reaction Mix and 5 μl of 4 mM DEVD-pNA (final concentration: 200 mM). The reaction solutions were incubated at 37°C for 1 h followed by measurement at OD 400 nm for colorimetric assay.

### Soluble TREM2 ELISA

96-well plate was coated overnight at 4 °C with TREM2 coating antibody (MAB17291-100, R&D Systems) in ELISA Coating Buffer (421701, Biolegend). It was then blocked in 25% Block Ace (BUF029, Bio-Rad laboratories) in PBS for 4 h at RT with constant shaking on a plate shaker. The plate was washed three times with wash buffer (0.1% Tween 20 in PBS) and incubated overnight at 4°C with plasma samples diluted 1:20 with 10% Block Ace in PBS. Plates were washed five times and incubated with mouse TREM2 biotinylated antibody (R&D Systems, BAF1729) for 2 h at RT with constant shaking. Plates were then washed and incubated with Streptavidin Poly-HRP40 Conjugate (65R-S104PHRP, Fitzgerald) for 1 h in the dark. After five additional washes the plates were developed by adding the TMB substrate (421501, Biolegend) and read at 620 nm.

### Bone marrow-derived macrophage (BMDM) culture

The femurs and tibias from C57BL/6J, Trem2 knock out, and DBA/2J mice were dissected, cleaned, and cut at the joints, and the bone marrow was flushed using a 25-gauge needle with RPMI 1640 with GlutaMAX™ Supplement (Gibco). Red blood cells were lysed using red blood cell lysis buffer (eBioscience). Cells were passed through a 100 μm filter. Cells were collected by centrifugation and washed multiple times with RPMI 1640. After counting, 10 million bone marrow cells were seeded per 10 cm non-treated tissue culture plates in RPMI 1640 Medium with 10% FBS (Omega Biosciences), 100 U/ml penicillin/streptomycin, 0.25 μg/ml Amphotericin B (Gibco), and 50 ng/ml M-CSF (Fujifilm Irvine Scientific Inc.). After 4 days of differentiation, 1/4 of additional fresh medium with same ingredients was added. After an additional 4 days of culture, cells were placed in a modular incubator chamber (Embrient, Inc) with 95% oxygen, 5% CO_2_ and were collected at different time points for downstream experiments.

### ChIP-seq library preparation

Chromatin immunoprecipitation (ChIP) was performed in biological replicates as previously described ^70^ with modifications as follows. Pulverized frozen mouse lung tissue powders or BMDMs were cross-linked with 2 mM disuccinimidyl glutarate (DSG) (ProteoChem) in PBS for 30 min at room temperature, and then directly a second crosslinking was performed by the addition of 1% (vol/vol) formaldehyde in PBS for 10 min. The crosslinking reaction was quenched by 0.125 M glycine (Sigma-Aldrich). The tissue powders or BMDMs were washed once with ice-cold PBS and pelleted at 1000 g for 5 min at 4°C. Crosslinked cells were resuspended in 100 ml of PIPA-NR buffer (20 mM Tris-HCl (pH 7.5), 150 mM NaCl, 1 mM EDTA, 0.5 mM EGTA, 0.4% sodium deoxycholate, 0.1% SDS, 1% NP-40, 0.5 mM DTT, 1 mM PMSF, 0.5 mM sodium butyrate, 1X protease inhibitor cocktail). Chromatin DNA was sonicated in a 96-well plate (Costar) using PIXUL™ Multi-Sample Sonicator (Active Motif) with the following setting: Process Time, 90 min; Pulse, 50 N; PRF, 1.00 kHz; Burst Rate, 20 Hz. Samples were centrifuged at 15000 rpm for 10 min at 4°C, and the supernatant was used for immunoprecipitation. 1% of the lysate was kept as ChIP input. The lysates were rotated with each antibody (Extended Data Table 2) pre-bound to 20 ml of beads (10 ml of Dynabeads protein A + 10 ml of Dynabeads protein G) overnight at 4°C. After the immunoprecipitation, beads were collected on a magnet and washed three times with PIPA-NR buffer and LiCl wash buffer (10 mM Tris-HCl (pH 7.5), 250 mM LiCl, 1% NP-40, 0.7% sodium deoxycholate, 1 mM EDTA, 1 mM PMSF, 0.5 mM sodium butyrate, 1X protease inhibitor cocktail), twice each with TET (10 mM Tris-HCl (pH 7.5), 1 mM EDTA, 0.2% Tween-20, 1 mM PMSF, 0.5 mM sodium butyrate, 1X protease inhibitor cocktail) and IDTE (10 mM Tris-HCl (pH 8.0), 0.1 mM EDTA, 0.05% Tween-20, 1 mM PMSF, 0.5 mM sodium butyrate, 1X protease inhibitor cocktail), and then resuspended in 25 μl of TT (10 mM Tris-HCl (pH 7.5), 0.05% Tween-20). The immunoprecipitated chromatin samples were used for library preparation with NEBNext Ultra II Library kit (NEB) according to the manufacturer’s instructions. DNA in 46.5 μl of NEB reactions was revers crosslinked by adding 4 μl of 10% SDS, 4.5 μl of 5 M NaCl, 3 μl of 500 mM EDTA, 1 μl of 20 mg/ml proteinase K and 20 µl of water by incubation for 1 h at 55°C, and then 30 min at 65°C. DNA was purified with 2 µl of SpeedBeads in 20% PEG 8000/1.5 M NaCl (Final 12% PEG 8000), and eluted with 25 μl of TT. The eluted DNA was amplified for 14 cycles in 50 μl of PCR reactions using NEBNext High-Fidelity 2X PCR Master Mix and 0.5 mM each of primers Solexa 1GA and Solexa 1GB. The amplified libraries were purified with 2 μl of SpeedBeads in 20% PEG 8000/2.5 M NaCl (Final 13% PEG 8000), eluted with 20 μl of TT, size selected using PAGE/TBE gel for 200-500 bp fragments by gel extraction, and pair-end sequenced on HiSeq 4000 or NovaSeq 6000 (Illumina) sequencer. ChIP input material (1% of sheared DNA) in 46.5 μl of TE was revers crosslinked by adding 4 μl of 10% SDS, 4.5 μl of 5 M NaCl, 3 μl of 500 mM EDTA, 1 μl of 20 mg/ml proteinase K and 20 µl of water by incubation for 1 h at 55°C, and then 30 min at 65°C. The input DNA was purified with SpeedBeads as described above and eluted with 25 μl of TT. The eluted input DNA was prepared for libraries and amplified as described for ChIP samples.

### RNA-seq library preparation

Total RNA in frozen lung tissue powders or BMDMs was purified using a Direct-zol RNA MicroPrep Kit (Zymo Reserch) as described by the manufacturer. mRNAs were enriched by incubation with Oligo d(T)_25_ Magnetic Beads (NEB). To fragment Poly A-enriched mRNA, mRNAs were incubated in 2X Superscript III first-strand buffer (Thermo Fisher Scientific) with 10 mM DTT at 94°C for 9 min. 10 μl of fragmented mRNAs were incubated with 0.5 μl of Random primers (3 μg/μl) (Thermo Fisher Scientific), 0.5 μl of Oligo dT primer (50 μM) (Thermo Fisher Scientific), 0.5 μl of SUPERase-In (Ambion) and 1 μl of dNTPs (10 mM) (Thermo Fisher Scientific) at 50°C for 1 min. After the incubation, 5.8 μl of water, 1 μl of DTT (100 mM), 0.1 μl of Actinomycin D (1 μg/μl) (Sigma-Aldrich), 0.2 μl of 1% Tween-20 and 0.5 μl of Superscript III (Thermo Fisher Scientific) was added and incubated on the following conditions: 25°C for 10 min, 50°C for 50 min, and 4°C hold. The mixture was purified with RNAClean XP beads (Beckman Coulter) as described by the manufacture and eluted with 10 μl of water. For second-strand synthesis with dUTP, the RNA/cDNA double-stranded hybrid was then added to 1.5 μl of Blue Buffer (Enzymatics), 1.1 μl of dUTP mix (10 mM dATP, dCTP, dGTP and 20 mM dUTP) (Promega), 0.2 μl of Rnase H (5 U/μl), 1.05 μl of water, 1 μl of DNA polymerase I (Enzymatics) and 0.15 μl of 1% Tween-20. The mixture was incubated at 16°C for 1 h. The resulting dUTP-marked dsDNA was purified with 2 μl of SpeedBeads in 20% PEG 8000/2.5 M NaCl (final 13% PEG 8000) and eluted with 40 μl of EB. The eluted dsDNA was end repaired by blunting, A-tailing and adapter ligation as previously described ^71^ using barcoded adapters (Bioo Scientific). The end repaired dsDNA was amplified for 15 cycles using 10 μL of NEBNext 2X High Fidelity PCR MM (NEB), and 0.2 μL each of primers Solexa 1GA and Solexa 1GB (stock concentration of 100 μM and final concentration of 1 μM). The amplified libraries were purified, and size selected with with 2 μl of SpeedBeads in 20% PEG 8000/2.5M NaCl (final 9% PEG 8000) and eluted with 12 μL of Elution Buffer, and pair-end sequenced on HiSeq 4000 or NovaSeq 6000.

### RNA-seq analysis

FASTQ files were processed to assess quality by determining general sequencing bias, clonality and adapter sequence contamination. RNA sequencing reads were aligned to the mm10 mouse reference genome using STAR ^72^. Gene expression levels were calculated using HOMER ^71^ by counting all strand specific reads within exons. Only the most abundant transcripts, including multiple alternative variants, were selected for each gene, and the genes with a length smaller than 250 bp were removed. Transcripts per million (TPM) were used to evaluate the correlation among replicates. Differential gene expression was calculated using DESeq2 ^73^ to assess both biological and technical variability between experiments. Unsupervised hierarchical clustering was used to cluster the gene expression in the heatmaps.

### ChIP-seq analysis

FASTQ files were mapped to the mm10 mouse reference genome with Bowtie2 ^74^. Peaks were called with HOMER (findPeaks) using parameters “-style factor -size 200”. Overlapping peaks from different conditions involved in a comparison were merged with HOMER’s mergePeaks tool (union) and annotated with HOMER’s annotatePeaks.pl. The raw reads of all replicate samples (Extended Data Table. 1) were quantified with HOMER (annotatePeaks.pl) using parameter “-raw”. DESeq2 was then used to identify differentially regulated peaks using an adjusted p-value cutoff of 0.1.

### Motif analysis

To identify motifs enriched in peak regions over the background, HOMER’s motif analysis “findMotifsGenome.pl” including known default motifs and *de novo* motifs was used ^71^. The background peaks used either from random genome sequences or from peaks in comparing condition were indicated throughout the main text and in the figure legends.

### Data visualization

ChIP-seq data were visualized by generating normalized bigwig read density files using HOMER and then viewed in the UCSC genome browser ^75^.

### Statistical analysis

The significance of differences in the experimental data were determined using GraphPad Prism 8.0 software. All data involving statistics are presented as mean ± SD or SEM. The number of replicates and the statistical test used are described in the figure legends.

## Acknowledgments

RNA-seq and ChIP-seq experiments were sequenced at the IGM Genomics Center, University of California, San Diego, La Jolla, CA. This publication includes data generated at the UC San Diego IGM Genomics Center utilizing an Illumina NovaSeq 6000 that was purchased with funding from a National Institutes of Health (NIH) SIG grant (#S10 OD026929). Tissue sections were processed at the UCSD Tissue Technology Shared Resource. The Tissue Technology Shared Resource is supported by a National Cancer Institute Cancer Center Support Grant (CCSG Grant P30CA23100). Y.A. was supported by the Japan Society for the Promotion of Science Overseas Research Fellowship (201860150). J.L.G. was supported by a NIH fellowship (F32-HL167318). We thank Chris Jansky for help with graphics.

## Author contributions

Y.A., N.J.S, C.K.G. and E.S. designed the project. Y.A., N.J.S., W.T., F.Z., J.L.C.R., C.S., S.J., M.M., R.H., K.C., D.D., S.G., J.L.G., D.M.L., M.T.L., C.B. and E.S. performed the experiments. C.K.G and E.S. secured the funding. Y.A., C.K.G. and E.S. wrote the manuscript.

## Competing interests

The authors declare no competing interests.

## Additional information

**Extended data** is available for this paper at OOO.

**Supplementary information** is available for this paper at OOO.

**Correspondence and requests for materials** should be addressed to E.S.

## Extended Data Figure legends

**Extended Data Fig. 1.**
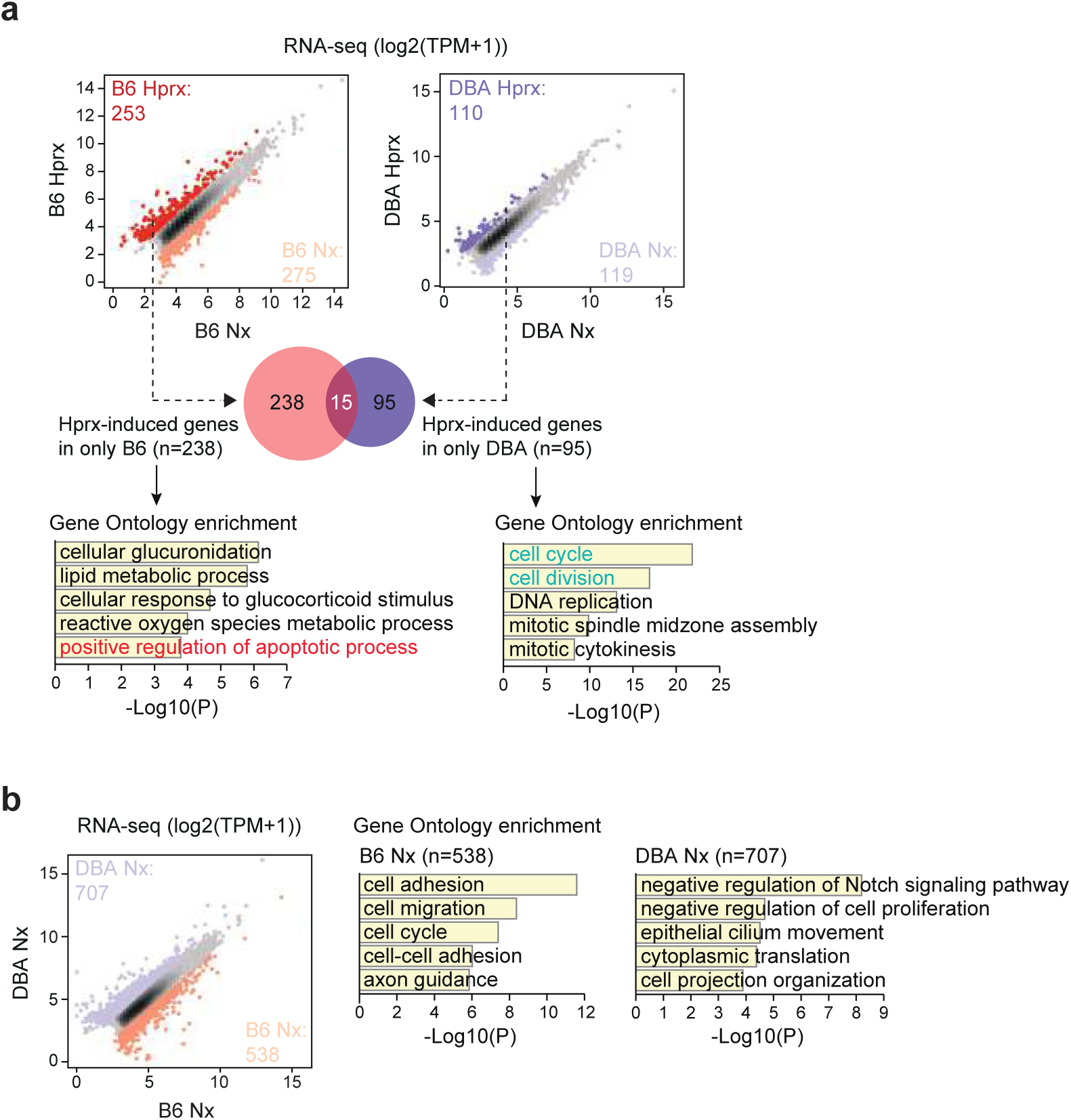
(a) Scatterplots of RNA-seq data in whole lung tissues showing differences in hyperoxia-induced genes in B6 and DBA mice. The left panel (same as Fig. 1e) shows the hyperoxia-regulated genes in B6 mice and right panel shows hyperoxia-regulated genes in DBA mice. Dark red dots in left panel: significant hyperoxia-induced genes in B6 mice, light red dots in left panel: significant hyperoxia-suppressed genes in B6 mice, dark purple dots in right panel: significant hyperoxia-induced genes in DBA mice, light purple dots in right panel: significant hyperoxia-suppressed genes in DBA mice (FC > 2; FDR < 0.05). Venn diagram shows the overlap between hyperoxia-induced genes in B6 (n = 253) and hyperoxia-induced genes in DBA mice (n = 110). Gene Ontology enrichment analysis of the 238 genes uniquely induced in B6 or 95 genes uniquely induced in DBA by hyperoxia. (b) Scatterplot of RNA-seq data in whole lung tissues showing baseline differences in gene expression between B6 and DBA mice (FC > 2; FDR < 0.05). Gene Ontology enrichment analysis of 538 genes enriched in the lungs of B6 mice or 707 genes enriched in the lungs of DBA mice.

**Extended Data Fig. 2.**
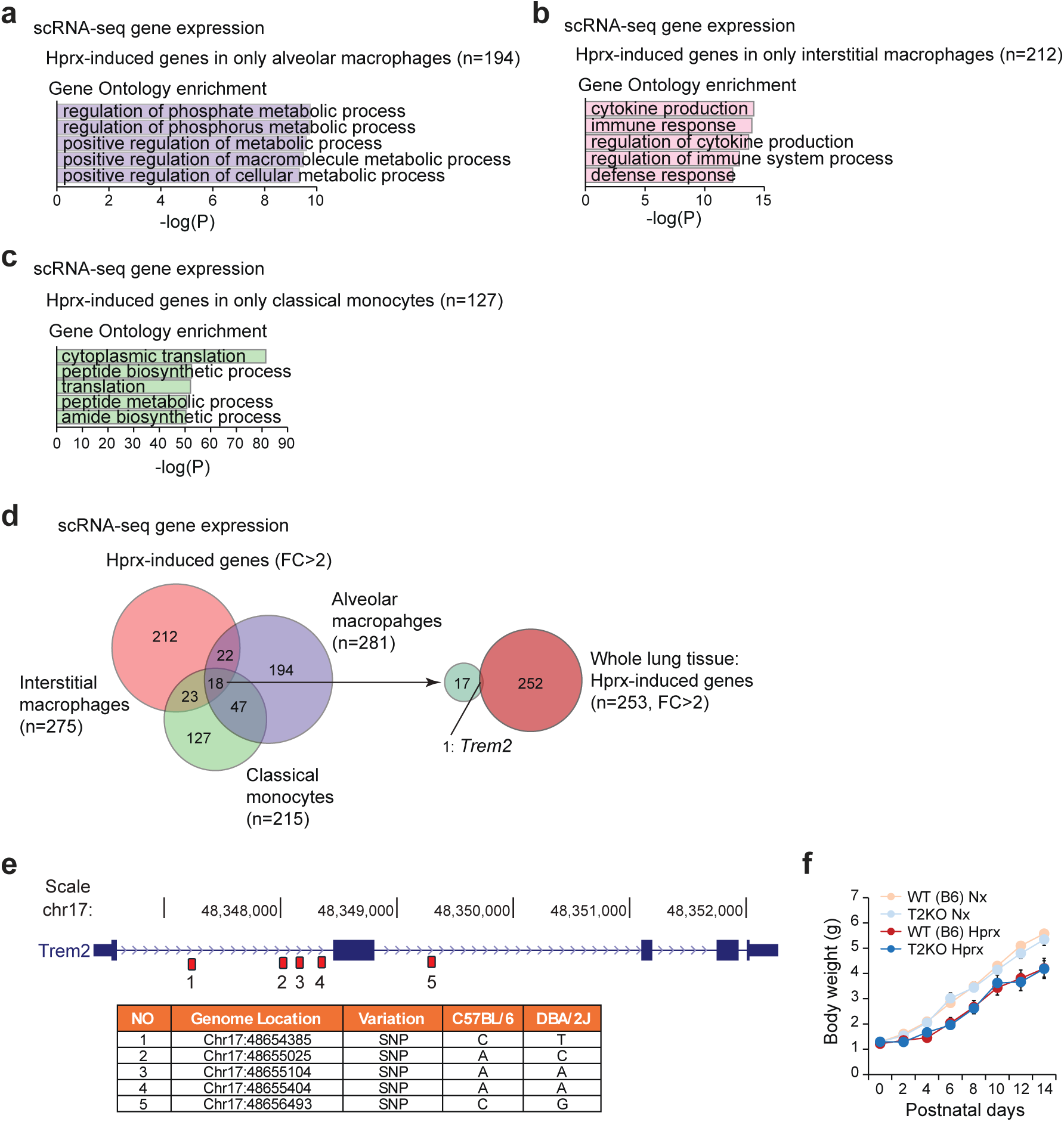
(a) Gene Ontology enrichment analysis of hyperoxia-induced genes specific for alveolar macrophages and not shared with interstitial macrophages and classical monocytes. Data reanalyzed from previously published scRNA-seq data ^1^. (b) Gene Ontology enrichment analysis of hyperoxia-induced genes specific for interstitial macrophages and not shared with alveolar macrophages and classical monocytes. Data reanalyzed from previously published scRNA-seq data ^1^. (c) Gene Ontology enrichment analysis of hyperoxia-induced genes specific for classical monocytes and not shared with interstitial macrophages and alveolar macrophages. Data reanalyzed from previously published scRNA-seq data ^1^. (d) Venn diagram showing the overlap between hyperoxia-induced genes (FC > 2; FDR < 0.05) in 3 different myeloid cell subsets; alveolar macrophages (n = 281), interstitial macrophages (n = 275), classical monocytes (n = 215) (left panel). Venn diagram showing the overlap between hyperoxia-induced genes in the three myeloid cell subsets (n = 18) and whole lung tissues (n = 253 in Fig. 1e) (right panel). (e) SNPs in the *Trem2* gene between B6 and DBA mice. (f) Body weight in hyperoxia-exposed WT (B6) (same samples as Fig. 1a) and T2KO mice. Data are mean ± SEM (n = 6 for each group).

**Extended Data Fig. 3.**
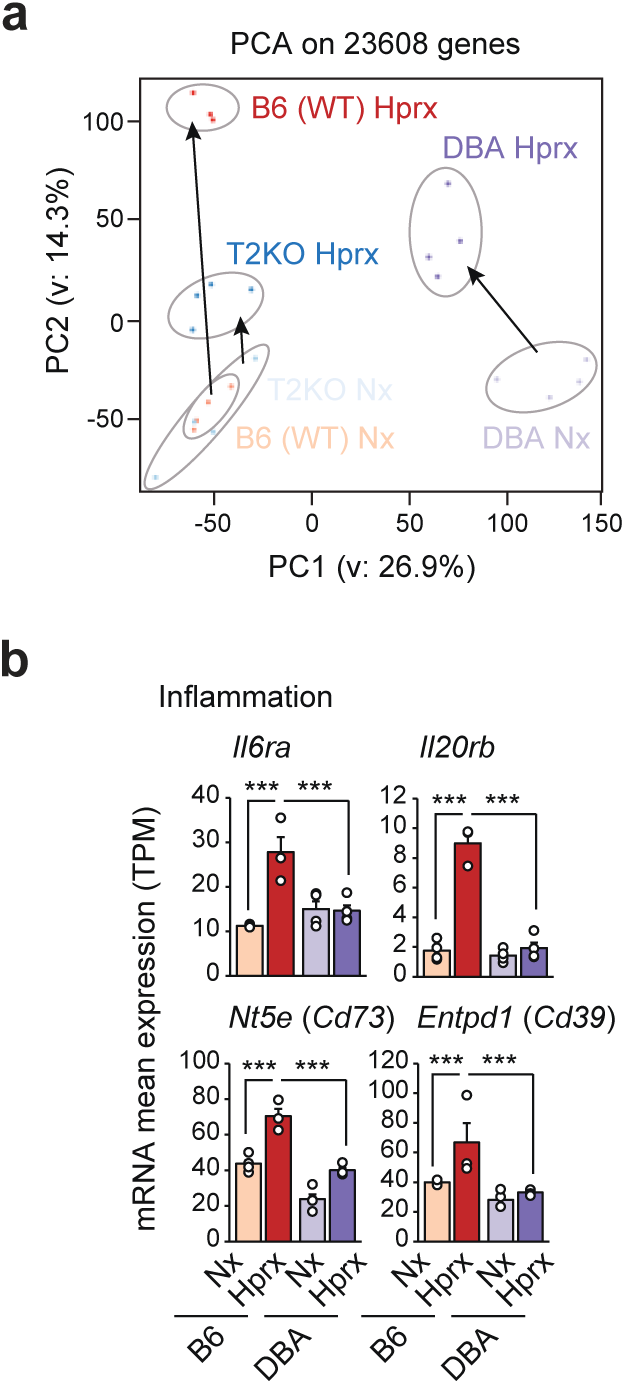
(a) Principal component analysis (PCA) of RNA-seq in whole lung tissues from B6 (WT), DBA and T2KO mice. (b) Bar plots for inflammatory gene expression in whole lung tissues between B6 and DBA mice after neonatal hyperoxia exposure. Data are mean ± SEM. DSeq2 was used for comparisons. *** *p*-adj < 0.001. Each dot represents one mouse.

**Extended Data Fig. 4.**
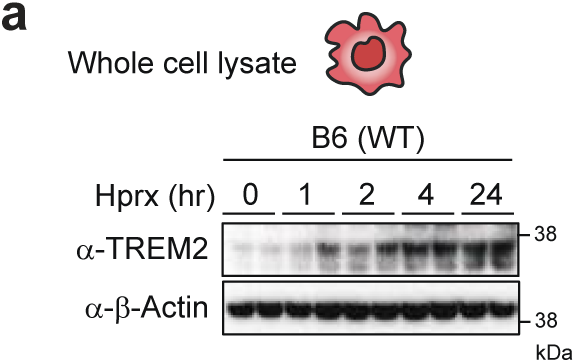
(a) Immunoblots of TREM2 protein in whole cell lysates of hyperoxia-exposed (1, 2, 4 or 24 h) BMDMs obtained from B6 (WT) mice and normoxic controls (0 h). Uncropped images of the blots are shown in Extended Data Fig. 7.

**Extended Data Fig. 5.**
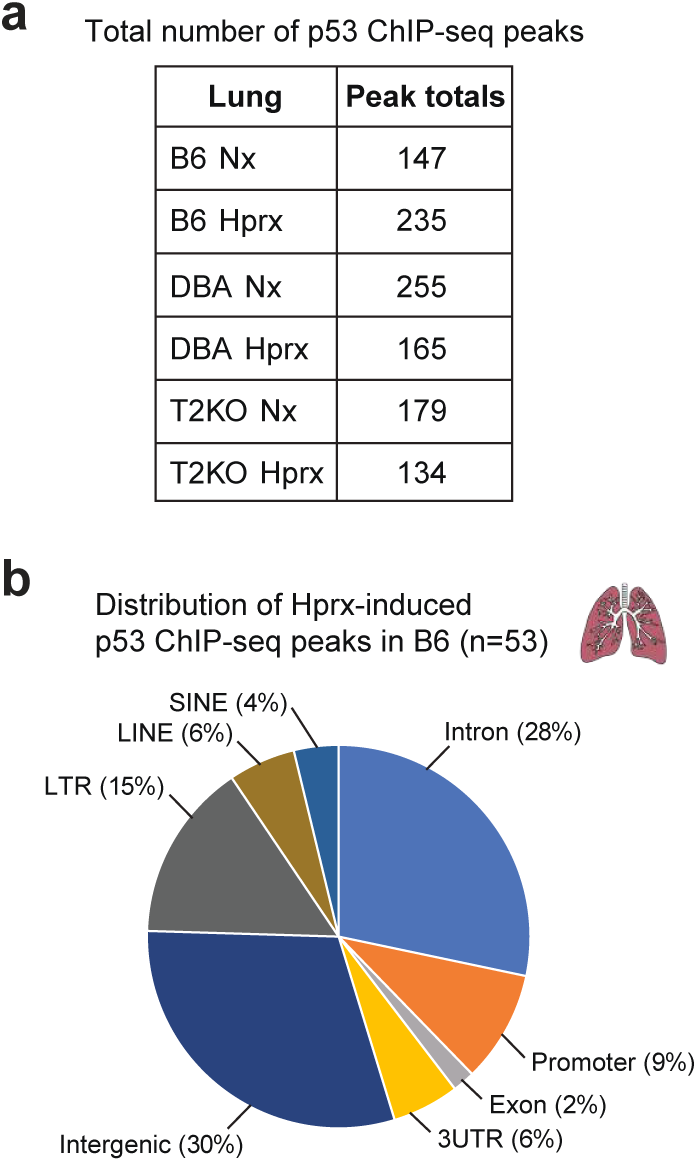
(a) Total number of p53 ChIP-seq peaks in whole lung tissues. (b) Distribution of hyperoxia-induced p53 ChIP-seq peaks (n=53) in whole lung tissues of B6 (WT) mice.

**Extended Data Fig. 6.**
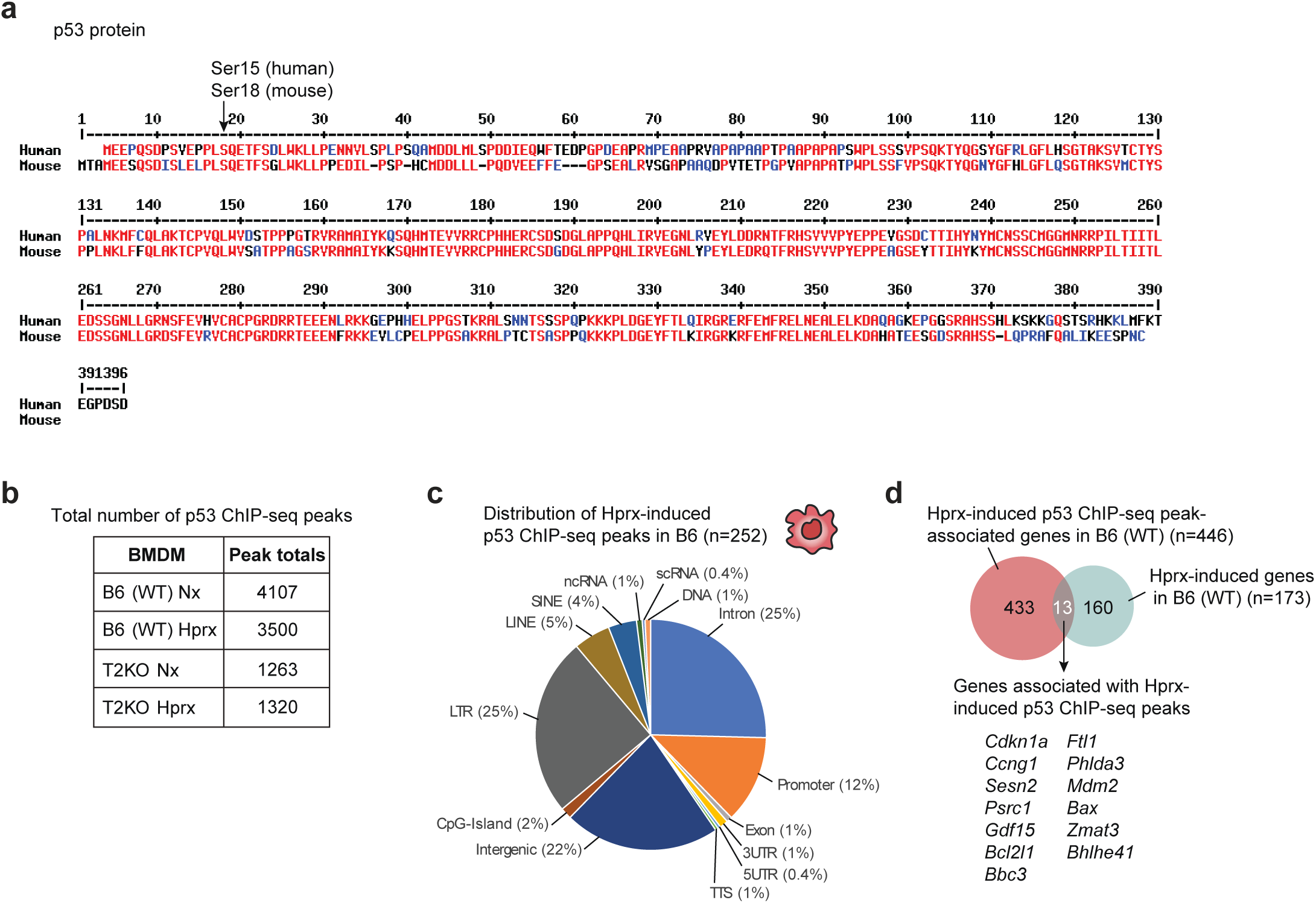
(a) Mouse and human p53 protein amino acid sequences. (b) Total number of p53 ChIP-seq peaks in BMDMs. (c) Distribution of hyperoxia-induced p53 ChIP-seq peaks (n=252) in BMDMs of B6 (WT) mice. (d) Venn diagram showing the intersection of hyperoxia-induced genes in BMDMs of B6 (WT) (n = 173, FC < 1.5; FDR < 0.05, Fig. 4d) and genes found near p53 ChIP-seq peaks (n = 446, 25 kbp from TSS) in BMDMs of B6 (WT) mice. *p* = 2.7E-10 (cumulative binomial distribution).

**Extended Data Fig. 7.**
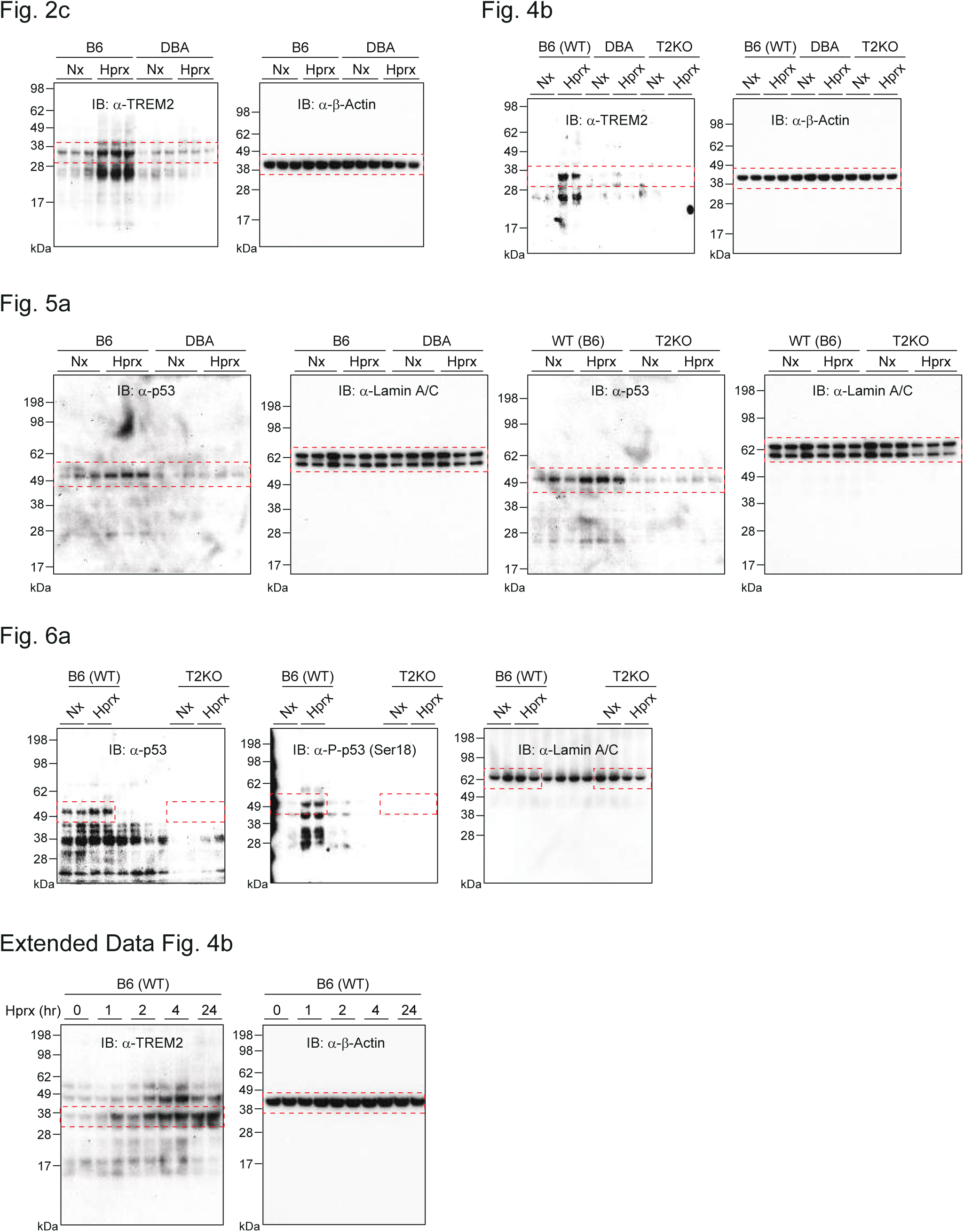
Uncropped immunoblot images for Fig. 2c, Fig. 4b, Fig. 5a, Fig. 6a and Extended Data Fig. 4b.

---

**Extended Data Table 1.**
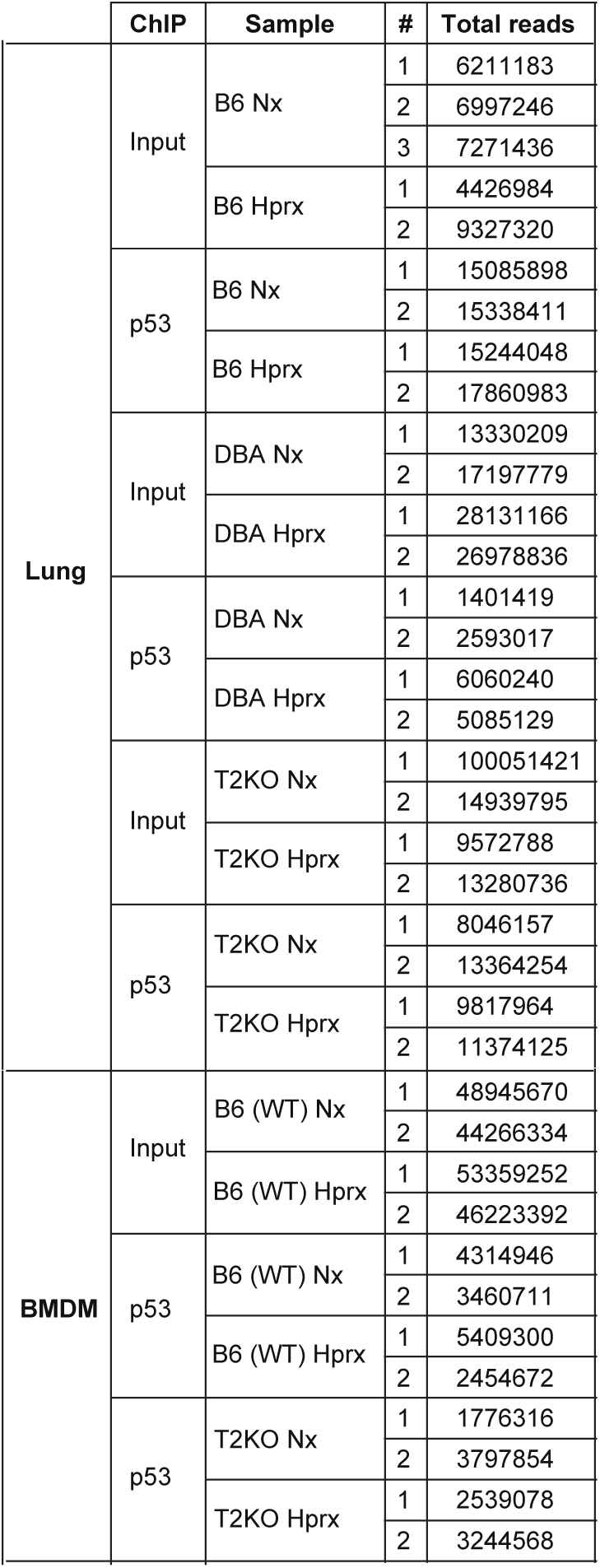
Total ChIP-seq reads included in analysis (Lungs and BMDMs)

**Extended Data Table 2.** Information of antibodies used in this study.

| Antibody |  | Source | Catalog No / Clone No | Dilutions / Concentrations |
| --- | --- | --- | --- | --- |
| anti-p53 | Monoclonal | Cell Signaling Technology | #2524S / 1C12 | 0.313 µg/ml for IB, 1.2 mg/assay for ChIP |
| anti-phospho-p53 | Polyclonal | Cell Signaling Technology | #9284T | 0.235 µg/ml for IB |
| anti-TREM2 | Monoclonal | R&D Systems | MAB17291-100 | 1 µg/ml for IB or ELISA |
|  | Monoclonal | R&D Systems | BAF1729 | 0.333 µg/ml for ELISA |
| anti-Lamin A/C | Polyclonal | Cell Signaling Technology | #2032S | 1:1000 for IB |
| anti-β-Actin | Monoclonal | Sigma-Aldrich | A2228 / AC-74 | 1:5000 for IB |

## Notes

### Competing Interest Statement

The authors have declared no competing interest.

